# Menopausal transition alters female skeletal muscle transcriptome

**DOI:** 10.1101/2023.11.27.568782

**Authors:** Hanna-Kaarina Juppi, Tia-Marje Korhonen, Tero Sievänen, Vuokko Kovanen, Urho M. Kujala, Pauliina Aukee, Neil Cronin, Sarianna Sipilä, Sira Karvinen, Eija K. Laakkonen

## Abstract

Menopause is associated with unfavorable changes in body composition. Skeletal muscle cells are targets of hormonal regulation and are affected by the interplay between coding and non-coding RNAs. Muscle transcriptome, including messenger-RNA (mRNA), long non-coding RNAs (lncRNAs) and microRNAs (miRNAs) has not previously been studied in women during the menopausal transition. Thus, we took a multi-RNA omics approach to get insight into transcriptome-wide events of menopause. Our study included baseline and follow-up muscle samples from seven early (EarlyMT) and 17 late perimenopausal (LateMT) women transitioning to early postmenopause during the study. Total RNA was sequenced and differential expression (DE) of the transcriptome was investigated. The potential gene functions were investigated with pathway analyses and protein level expression with Western Blot. We found 30 DE mRNA genes in EarlyMT and 19 in LateMT participating in pathways controlling cell death, growth, and interactions with external environment. Lack of protein level changes may indicate a specific role of the regulatory RNAs during menopause. Ten DE lncRNA transcripts were identified but did not result in DE lncRNAs genes. No DE miRNAs were found. Despite the lack of DE findings in regulatory RNAs, we identified putative regulatory networks, likely to be affected by estradiol availability. The changes in gene expression were correlated with observed changes in body composition variables, indicating muscularity and adiposity regulators to be affected by menopausal transition. In essence, the observed DE genes and their regulatory networks may offer novel mechanistic insights on factors affecting body composition during and after menopause.

## Introduction

Approximately half of the human population confronts menopause, which leads to a multitude of physiological changes and is associated with increased disease risks. Skeletal muscle is among the affected tissues, but little is known about the underlying transcriptome-wide changes occurring during menopausal transition. Skeletal muscle, which comprises approximately 40 percent of the total body weight, is not only responsible for movement, balance, heat production and amino acid storage but also has an important role in metabolism and inter-tissue signaling (Frontera & Ochala, 2015). Muscle mass and function are highly determined by the proteins synthesized and degraded. While myostatin, mTOR and MyoD are examples of well-established protein-level contributors to muscle mass and metabolism (Bentzinger et al., 2012; Bodine et al., 2001), the RNA-level regulatory network is still imperfectly characterized especially in women. Due to the important metabolic role of skeletal muscle, various aspects of RNA signaling within the tissue have also the potential to contribute to total body health and metabolism (Martone et al., 2020).

Discovery of the non-coding RNAs (ncRNAs), such as long non-coding RNAs (lncRNAs) and microRNAs (miRNAs) has led to a widened perspective on the regulation of gene expression. LncRNAs are classified as ≥200 nucleotide-long oligonucleotides lacking protein-coding potential (Statello et al., 2021). LncRNAs control gene expression through pre-transcriptional and post-transcriptional mechanisms such as regulation of chromatin accessibility and transcription factor recruitment and by affecting messenger RNA (mRNA) stability or competing with miRNAs on target mRNAs (Fernandes et al., 2019). miRNAs are short ncRNA molecules known to inhibit protein translation of about 60% of mRNA genes (Friedman et al., 2009). To date, more than 170,000 different lncRNA transcripts (Zhao et al., 2020) and approximately 2600 different mature miRNA sequences have been identified from the human genome (*miRBase: The microRNA Database*, 2022). As different RNA species have significant regulatory roles on each other, understanding more of the complicated functional RNA signaling networks in tissues is critical to understand how they orchestrate human physiology and health.

Natural menopause at midlife is a time of dramatic hormonal changes caused by aging-related alterations in the ovaries. This results in permanently lowered female sex hormone levels, including progesterone (P4) and the biologically most potent estrogen, estradiol (E2). While E2 is known to have beneficial effects in several tissues, including skeletal muscle, the executive mechanisms have remained largely unknown. Higher systemic E2 levels and pre- or perimenopausal status have been associated with higher muscle mass (Juppi et al., 2020; Sowers et al., 2007), better muscle quality and strength (Bondarev et al., 2018; Sipilä et al., 2001), lower body fat mass (Greendale et al., 2019; Juppi et al., 2022) and lower risk for metabolic syndrome (Janssen et al., 2008). With respect to muscle transcriptome, only a few cross-sectional studies examining the potential role of E2 and other menopause-related hormones on muscle properties have thus far been conducted. In postmenopausal women, E2 levels have been shown to associate with differences in the part of the muscle mRNA transcriptome that controls muscle mass, performance and metabolism, indicating E2-mediated regulation (Pöllänen et al., 2007; Ronkainen et al., 2010). In the case of small ncRNAs, E2 has been shown to regulate several growth, autophagy and glucose metabolism-linked miRNAs in human skeletal muscle (Olivieri et al., 2014), but to our knowledge, estrogenic regulation of lncRNAs has not been previously studied in human skeletal muscle. The few in vivo studies available suggest a regulatory role of female sex hormone levels in muscle lncRNA expression (Chai et al., 2019; J. Wang et al., 2017).

While skeletal muscle significantly impacts human health, and menopause is known to negatively impact skeletal muscle, our understanding of the underlying mechanisms, such as how menopausal transition impacts skeletal muscle transcriptome, has remained limited. The aim of this study was to investigate the longitudinal associations of the menopausal transition with the expression of mRNA, lncRNA and miRNA in human skeletal muscle. We also investigated whether changes in the transcriptome were associated with body composition variables.

## Results

### Characteristics of the study participants: EarlyMT and LateMT groups differed in the rate of decrease of estradiol

The participating women were assigned into early (EarlyMT, n = 7) or late (LateMT, n = 17) menopausal transition groups based on their baseline menopausal status being early or late perimenopausal, respectively. During an individualized follow-up period, each participant’s menopausal transition was monitored by checking their follicle-stimulating hormone (FSH) levels and menstrual diaries every three to six months until they were regarded as early postmenopausal(Kovanen et al., 2018). The mean follow-up time was 1.5 ± 0.9 years for the EarlyMT and 1.0 ± 0.6 years for the LateMT group. At baseline, women in the EarlyMT group were on average 52.6 ± 2.5 years old and in the LateMT group 52.2 ± 2.0 years old. In line with our earlier findings using a larger sample of this same study cohort(Juppi et al., 2020, 2022), lean body mass tended to decrease and fat mass to increase (**Supplementary Table 1**). A typical menopausal transition-associated hormonal change was observed in both groups, i.e., FHS levels increased and E2 decreased although the E2 decrement was significantly greater in the EarlyMT than in the LateMT group (p = 0.034) (**Supplementary Table 1**).

### Over 18,000 mRNA genes were expressed in muscle including structural proteins, hormone receptors and steroidogenic enzymes

Preprocessing of the sequencing data was done separately for sequences representing long (mRNAs + lncRNAs) and short (miRNAs) RNA classes (**Figure 1a**). As well known, transcription of a single gene can result in several splice variants and all of them can be present as single RNA transcripts in the sequencing data of the long RNA. Here we investigated long RNA expression considering both the transcript and the gene levels. In the latter, transcripts resulting from the same gene were combined by *tximport* (Soneson et al., 2016). For simplicity and clarity, we use the terms transcript and gene to distinguish if the observation or analysis refers to the transcript or gene level.

**Figure 1.**
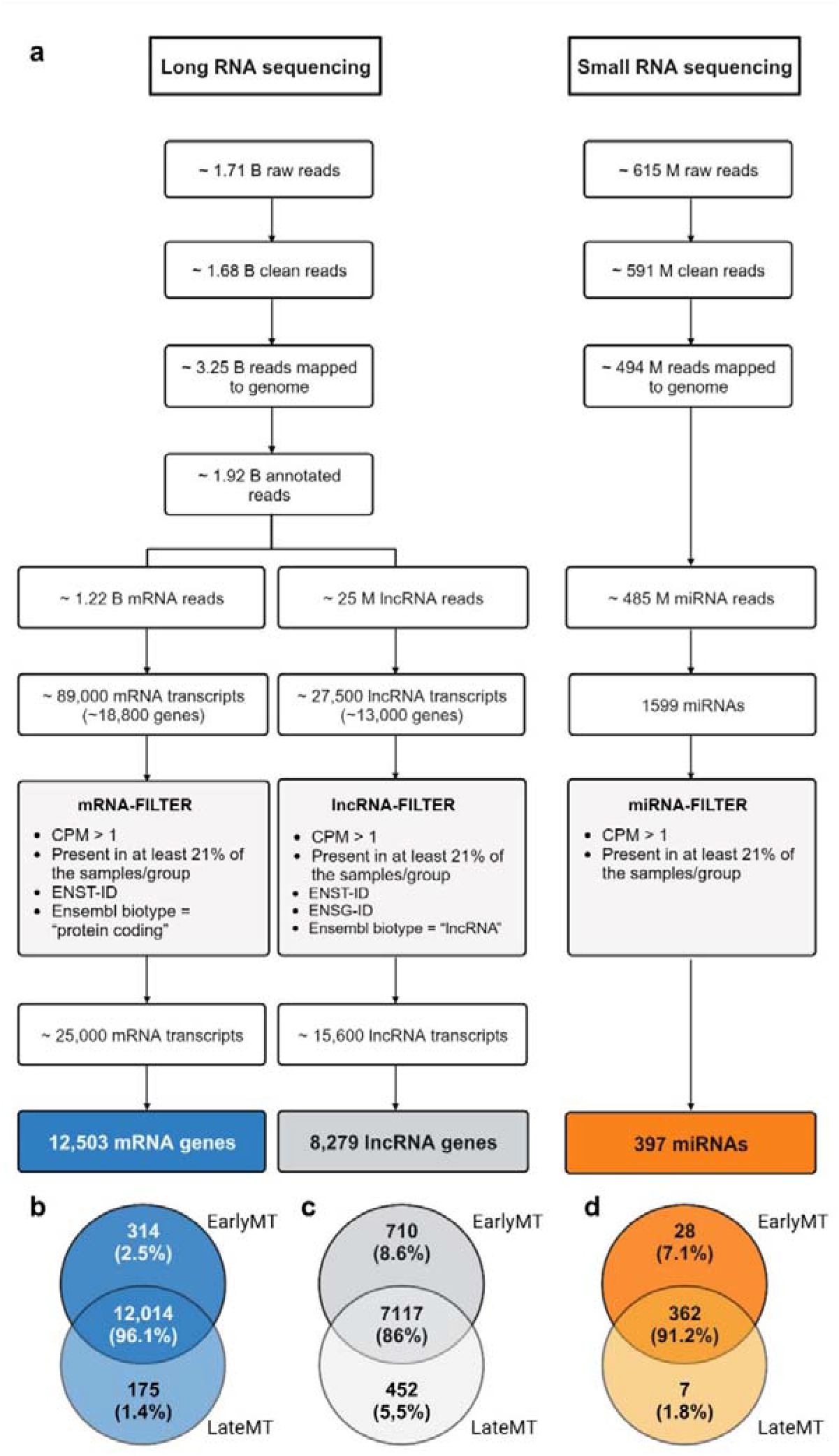
Preprocessing and analytical flow of the sequencing data. **a** Flowchart of the data inclusion process. Venn-diagram presenting **s**hared and unshared **b)** mRNA genes, **c)** lncRNA genes and **d)** miRNAs in EarlyMT and LateMT. Abbreviations: B, billion; M, million; mRNA, messenger RNA; lncRNA, long non-coding RNA; miRNA, microRNA; CPM, counts per million; ENST-ID, Ensembl Transcript ID that is a unique identifier for each transcript of a gene; ENSG-ID, Ensembl Gene ID that is a unique identifier for each gene; EarlyMT, a group of women transitioning from early perimenopause to early postmenopause; LateMT, a group of women transitioning from late perimenopause to early postmenopause

Of the long RNA clean reads, 96% were mapped to the reference genome. Approximately 89,000 long RNA transcripts were classified as mRNA representing 18,769 different genes (**Figure 1a**). The most abundant mRNA transcripts included several splice variants of typical skeletal muscle structural proteins such as *MYH7*, *MYH2*, *ACTA1, TTN* and *NEB*. After filtering, a total of 12,503 different mRNA genes remained (for filter specifications, see **Figure 1a** and methods). Due to the study aims, we had a special interest in inspecting gene expression levels of sex hormone receptors, steroidogenic enzymes, and proteins related to muscle function and metabolism (**Supplementary Table 2**). The receptor for FSH (*FSHR*) and steroidogenic enzymes *HSD3B2*, *HSD17B2* and *CYP19A1* were also detected among the sequences data but their count numbers were too low to meet the criteria to be classified as an expressed transcript or gene and were thus omitted from further analyses. Of the included mRNA genes, 314 were expressed only in EarlyMT and 175 only in LateMT (**Figure 1b** and **Supplementary Table 3**).

### Overrepresentation analysis of the mRNA genes pointed towards changes in extracellular matrix

To get an insight into the potential functional differences in the expressed mRNA genes between EarlyMT and LateMT groups, we did a focused gene ontology (GO) analysis of the expressed mRNA genes that were not shared between EarlyMT and LateMT (**Figure 1b**). The 314 EarlyMT-specific genes were enriched into 35 GO terms (adjusted p-value (p_adj_) < 0.05) while no statistically significant enrichment was observed for the 175 LateMT genes. The EarlyMT-specific enriched genes were connected (**Supplementary Table 4**), for example, to the biological processes related to immune and inflammatory responses (15 GO pathways) and metabolism nucleic acids and other molecules (15 GO pathways).

Next, we wanted to get overall insight on the changes in active pathways during menopausal transition. Therefore, we did Gene Set Enrichment Analysis (GSEA) by using all the prefiltered mRNA genes (**Figure 1a**) expressed in Earliest and LateMT as input data to be compared against Reactome and GO biological processes databases. The most significant findings are shown in **Table 1**. As a generalization of the GSEA results, we conclude that in both EarlyMT and LateMT the mRNA genes were enriched in cellular processes controlling the cell survival and interactions with the external environment.

**Table 1.** Significant results (p_adj_ <0.1) from GSEA analysis of the protein-coding genes.

| EarlyMT |  |  | LateMT |  |  |
| --- | --- | --- | --- | --- | --- |
| | Hits/Gene<br>set size | $p_{adj}$ | | Hits/Gene<br>set Size | $p_{adj}$ |
| **<u>Gene Ontology: Biological Process</u>** |  |  |  |  |  |
| External encapsulating structure organization | 103/210 | 0.005 | Cell substrate adhesion | 80/275 | 0.082 |
| Cell substrate adhesion | 83/277 | 0.065 | External encapsulating structure organization | 72/210 | 0.082 |
| Receptor mediated endocytosis | 67/183 | 0.065 |  |  |  |
| Collagen biosynthetic process | 18/30 | 0.065 |  |  |  |
| Regulation of cell substrate adhesion | 36/163 | 0.097 |  |  |  |
| **<u>Reactome</u>** |  |  |  |  |  |
| ECM organization | 95/203 | 0.002 | ECM organization | 71/202 | 0.001 |
| Interferon signaling | 57/151 | 0.084 | Syndecan interactions | 11/24 | 0.046 |
| Degradation of the ECM | 43/83 | 0.084 | ECM proteoglycans | 18/51 | 0.064 |
| Regulation of insulin-like growth factor transport and uptake by insulin-like growth factor binding proteins | 29/74 | 0.084 |  |  |  |

EarlyMT, a group of women transitioning from early perimenopause to early postmenopause; LateMT, a group of women transitioning from late perimenopause to early postmenopause; ECM, extracellular matrix; GSEA, gene set enrichment analysis; *Hits* refer to genes found in our dataset, while *Gene set size* refers to the gene sets used on the database.

### Human muscle expresses nearly 13,000 different lncRNA genes

Altogether ∼27,500 long RNA transcripts constituting 12,942 different genes were classified as lncRNAs (**Figure 1a**). The most abundant lncRNA transcripts included *MALAT1*, *NORAD*, *NEAT1* and two miRNA host genes (for miR-1 and miR-3648). After filtering, a total of 8279 different lncRNA genes were left for further analysis. Of these, 710 were only expressed in EarlyMT while 452 were unique to LateMT (**Figure 1c** and **Supplementary Table 5**). Of the 20 topmost expressed genes, 19 were common in both groups and included *NORAD*, *MALAT1*, *XIST*, *NEAT1*, *MIR1-1HG*, *H19*, *SNHG14*, *KCNQ1OT1*, *FGD5-AS1*, *SNHG5*, *SNHG16*, *MIR133A1HG*, *MIR193BHG*, *LINC01405*, *ZNF710-AS1*, *NUTM2A-AS1* and five other lncRNA genes with currently uncharacterized function. *ZNF710-AS* was only included in the EarlyMT top 20 list, and *NUTM2A-AS1* only occurred in the LateMT top 20 list.

### The top 20 miRNAs constituted the majority of all observed miRNA reads

Reference genome alignment of the clean small RNA reads was on average 83% (**Figure 1a**). Altogether 1599 different miRNAs were sequenced. After filtering, 397 miRNAs were included in the further analysis. Of these, 362 were shared by both groups, while 28 were only expressed in EarlyMT and seven only in LateMT (**Figure 1d** and **Supplementary Table 6**). The 20 most abundant miRNAs in both groups constituted 97.2 % of all miRNA reads and were with two exceptions the same in both groups including let-7a-5p, 7f-5p, -7g-5p, -7i-5p, and miR-1-3-p, -21-5p (only present in EarlyMT top 20 list), -26a-5p, -27b-3p, - 30a-5p, -30d-5p, -99a-5p, -126-3p, -133a-3p, -143-3p, -148a-3p, -206, -378a-3p, -378c, -378d, -451a, and -486-5p (only present in LateMT top 20 list). Notably, of the muscle-enriched myomiRs (Abkhooie et al., 2021), four (miR-1-3-p, -133a-3p, -206 and -486-5p) were also present in the top 20 list.

### Menopausal transition is associated with changes in mRNA gene expression

We found 30 mRNA genes to be differentially expressed (DE) within EarlyMT and 19 within LateMT (for all p_adj_ < 0.05, log fold change [LFC] ≥ ± 1.5, further details in **Table 2** and **Supplementary** Figure 1). EarlyMT DE genes included for example structural protein *ELN*, steroidogenic enzyme *SRD5A1* and apoptosis linked *PIDD1*. LateMT DE genes included e.g., molecular switch regulator *NUDT4*, extracellular matrix component *ECM1*, and the negative regulators of apoptosis *NAA35* and *BIRC6*. In both groups, the DE genes included several regulators of transcription, such as *GTF2F2*, *E2F3*, *MAFK, ZEB1, KIAA0355/GARRE*, and zinc fingers *ZNF84* and *ZNF611*. The DE genes also included *MKNK1*, *JAK2* and *MYD88*, which are components of the important growth, metabolism and adaptation regulating MAPK, mTOR and PI3K/Akt signaling pathways in the muscle.

**Table 2.**
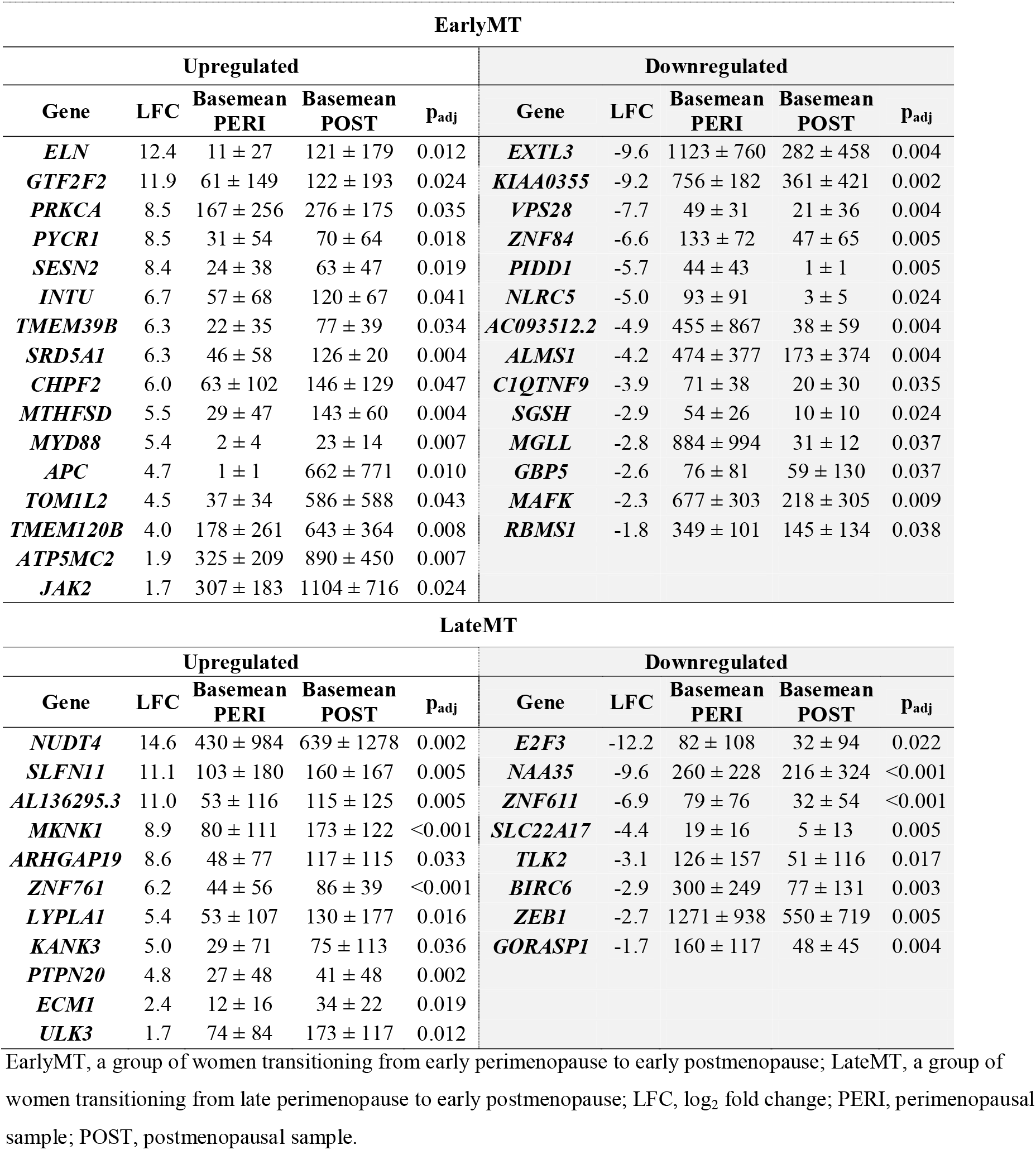
Differentially expressed protein-coding genes (POST vs. PERI) in the EarlyMT and LateMT groups.

No DE lncRNA genes or miRNAs were found. However, at the individual transcript level, we found in total ten DE lncRNA transcripts in the EarlyMT and LateMT groups (p_adj_ < 0.05 and LFC ≥ ± 1.5, **Supplementary Table 7**). In EarlyMT, *OSER1-DT*, *MALAT1* and *AC025171.1* were downregulated. In LateMT, *BAIAP2-DT* and *LINC02541* were upregulated and *AC083798.2*, *AL050309.1*, *LINC00667*, *IQCH-AS1* and *ENTPD1-AS1* were downregulated between the post- and perimenopausal states.

### Correlation network and Ingenuity Pathway Analysis predicted associations between menopause-related hormones, transcriptome, and downstream functions

To inspect relationships between the three RNA classes and to identify the most significant associations, we used sparse partial least squares-discriminant analysis (sPLS-DA) in *mixOmics* (Rohart et al., 2017). sPLS-DA reduces the dimensionality of the multi-omics data (here mRNA, lncRNA and miRNA expression data) by selecting the most predictive or discriminant features (Lê Cao et al., 2011). After applying a correlation threshold r ≥ 0.7, all remaining correlations between the identified key features were positive. The results were then replotted with Cytoscape and presented in **Figure 2**. Two distinct correlation networks were observed: one around mRNA genes *ALMS1* and *RBMS1*, and the other one around mRNA genes *SGSH* and *KIAA0355*. MyomiRs miR-486, miR-133a and miR-133b were part of these networks. Although the performed analysis does not allow directional causal conclusions to be made, it is tempting to speculate that involved regulatory RNAs (lncRNAs and miRNAs) would form a regulatory network for controlling the expression of *ALMS1*, *RBMS1*, *SGSH* and *KIAA0355*.

**Figure 2.**
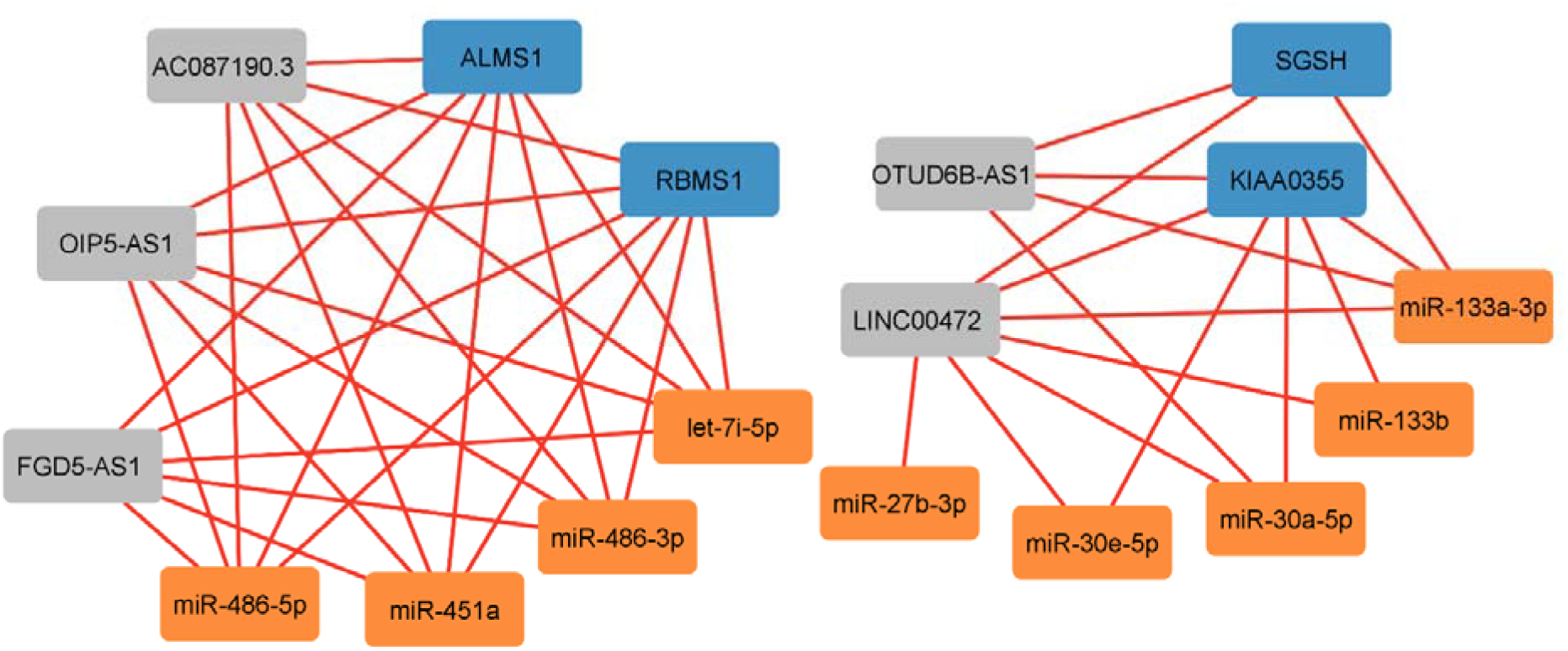
The multi-RNA correlation networks resulting from multiblock-analysis of the observed longitudinal changes in the expression among 24 women transitioning from early or late perimenopause to early postmenopause. Only correlations > 0.7 are shown. Blue = mRNA, grey = lncRNA, orange = miRNAs.

Using Ingenuity Pathway Analysis (IPA) we further explored in which cellular pathways the DE mRNA genes were enriched. The advantage of IPA is that it applies the researcher-input expression data and its own curated knowledge base to compute z-scores with the predictive value of potential activation (positive) or inhibition (negative) of pathways or none indicating inconsistencies between expression data and knowledge base (Krämer et al., 2014). Analyses were done separately for the EarlyMT and LateMT groups. We present the top 15 IPA-identified muscle-related enriched (p<0.05) canonical pathways in **Figure 3a** and **3b**. For LateMT group, IPA was not able to provide prediction towards pathway activation or inhibition for any of the pathways. The total list of pathways and their molecules can be found in **Supplementary Tables 8 and** 9. Interestingly, the identified pathways included several pathways related to hormonal regulation, e.g., androgen signaling, estrogen receptor signaling and gonadotropin-releasing hormone (GnRH) signaling, as well as regulation of cell death and energy metabolism.

**Figure 3.**
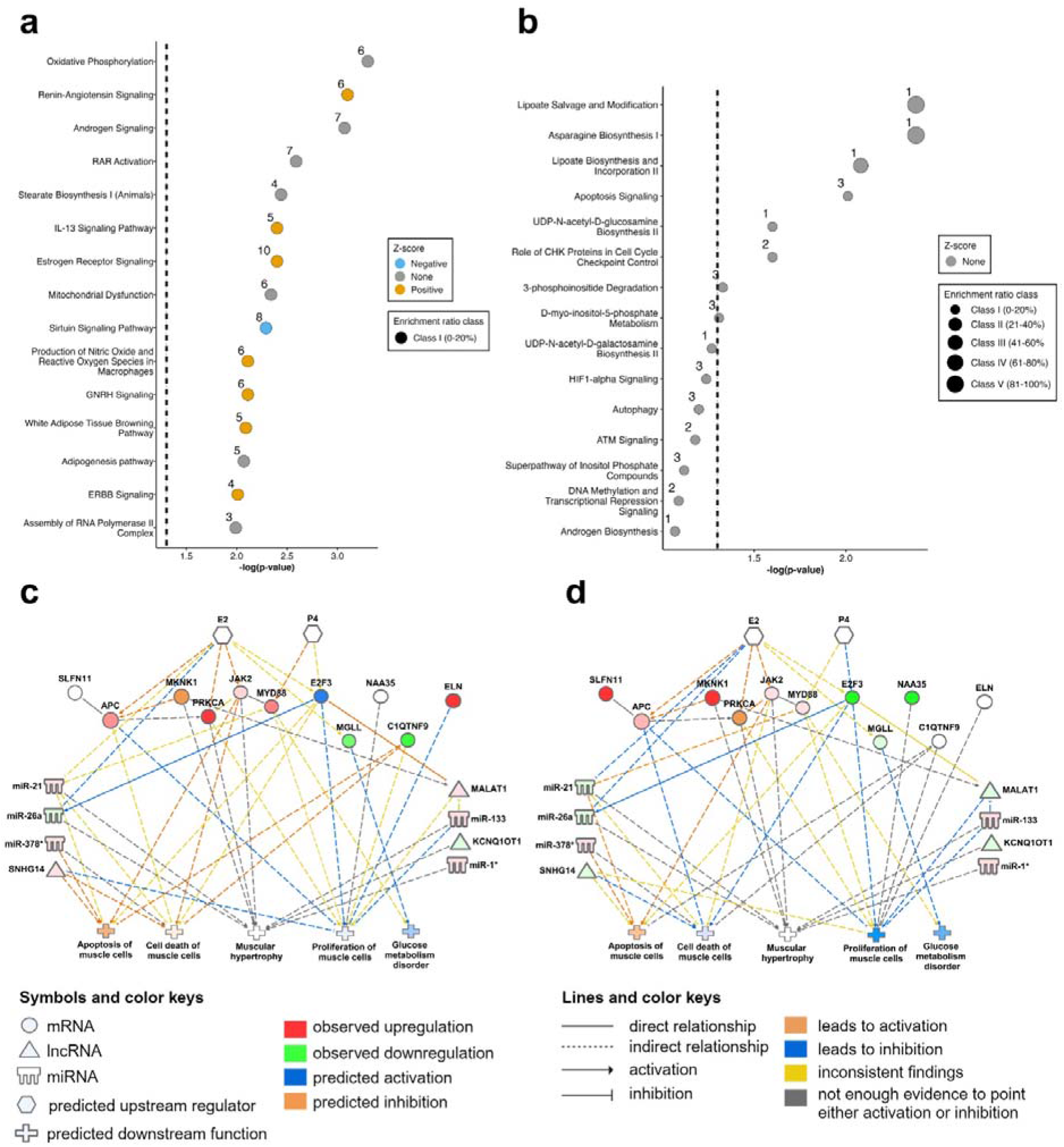
Results of the Ingenuity Pathway Analysis. **a** Top 15 most significant canonical pathways in EarlyMT and **b** LateMT groups. Only muscle-relevant pathways are shown. In a and b, circle size refers to the ratio class of observed genes to all genes in the pathway. The z-score of the pathway is used to predict the activation (positive, orange) or inhibition (negative, blue) of the pathway. If neither could be predicted, circle color was left grey.The number on the left side of the circle refers to the actual number of the observed genes. The dashed line indicates significance (log (p-value) > 1.3). **c** Results of the My Pathway analysis to identify potential upstream regulators and the affected downstream functions focusing on **c** EarlyMT and **d** LateMT groups. For both analyses, the same list of differentially expressed mRNA genes and top 20 expressed lncRNA and miRNAs were used as input while color keys represent observations from originating from **a** EarlyMT and **b** LateMT Abbreviations: EarlyMT, a group of women transitioning from early perimenopause to early postmenopause; LateMT, a group of women transitioning from late perimenopause to early postmenopause

Next, we wanted to inspect, by using the My Pathway analysis option in IPA, if E2, P4 or FSH as the major female hormones subjected to change during the menopausal transition, could function as upstream regulators of the DE genes thereby affecting the muscle tissue decrements observed at the phenotype level. The DE mRNA genes from **Table 2** and the top 20 expressed regulatory RNAs were included in the My Pathway analysis. **Figures 3c** and **d** show the results of the combined analysis, where all DE mRNA genes that were found to associate either with upstream regulators or downstream functions are shown in the same network despite being differentially expressed only in either EarlyMT or LateMT group. This presentation provides a scene over the whole menopausal transition period while the colors represent the transition from early (**3c**) or late (**3d**) perimenopause to early postmenopause. FSH was not found to regulate any of the observed DE genes, thus it is not included in the figures. E2 was predicted to be an upstream regulator of *APC*, *PRKCA*, *JAK2, E2F3* and *MGLL*, while P4 was predicted to regulate *MYD88* and *E2F3*. E2- and P4-regulated genes were found to be associated with several aspects of muscle function and morphology including muscle cell apoptosis, death and proliferation, muscle hypertrophy, and glucose metabolism disorder. Steroidogenesis and satellite cell function also appeared as associate pathways but for the sake of clarity of the figures, are not shown. In EarlyMT, muscle cell apoptosis and cell death were predicted to be activated, while glucose metabolism disorder and muscle cell proliferation were predicted to be inhibited (**Figure 3c**). Using expression data from LateMT, muscle cell proliferation, cell death and glucose metabolism disorder were predicted to be inhibited, while muscle cell apoptosis was expected to be activated (**Figure 3d**). IPA also predicted, that within the included top 20 regulatory RNAs there were several downstream targets of estrogenic regulation as well as associations between the muscle tissue properties and regulatory RNAs.

### Protein level analyses suggest regulatory steps before translation

We further wanted to validate whether some of the mRNA level differences of muscle regeneration and metabolism regulators such as ZEB1, MYD88, PRKCA, JAK2, E2F3 and APC were directly translated to protein expression level. Unfortunately, the antibodies for JAK2, E2F3 and APC turned out to be unspecific. Western blot analyses of paired samples revealed no difference in protein levels for ZEB1 or PRKCA in either of the menopausal groups, while there was a trend for upregulation of MYD88 in LateMT (p = 0.070) (**Figure 4**), similar to what was seen at the gene level in EarlyMT (**Table 2**).

**Figure 4.**
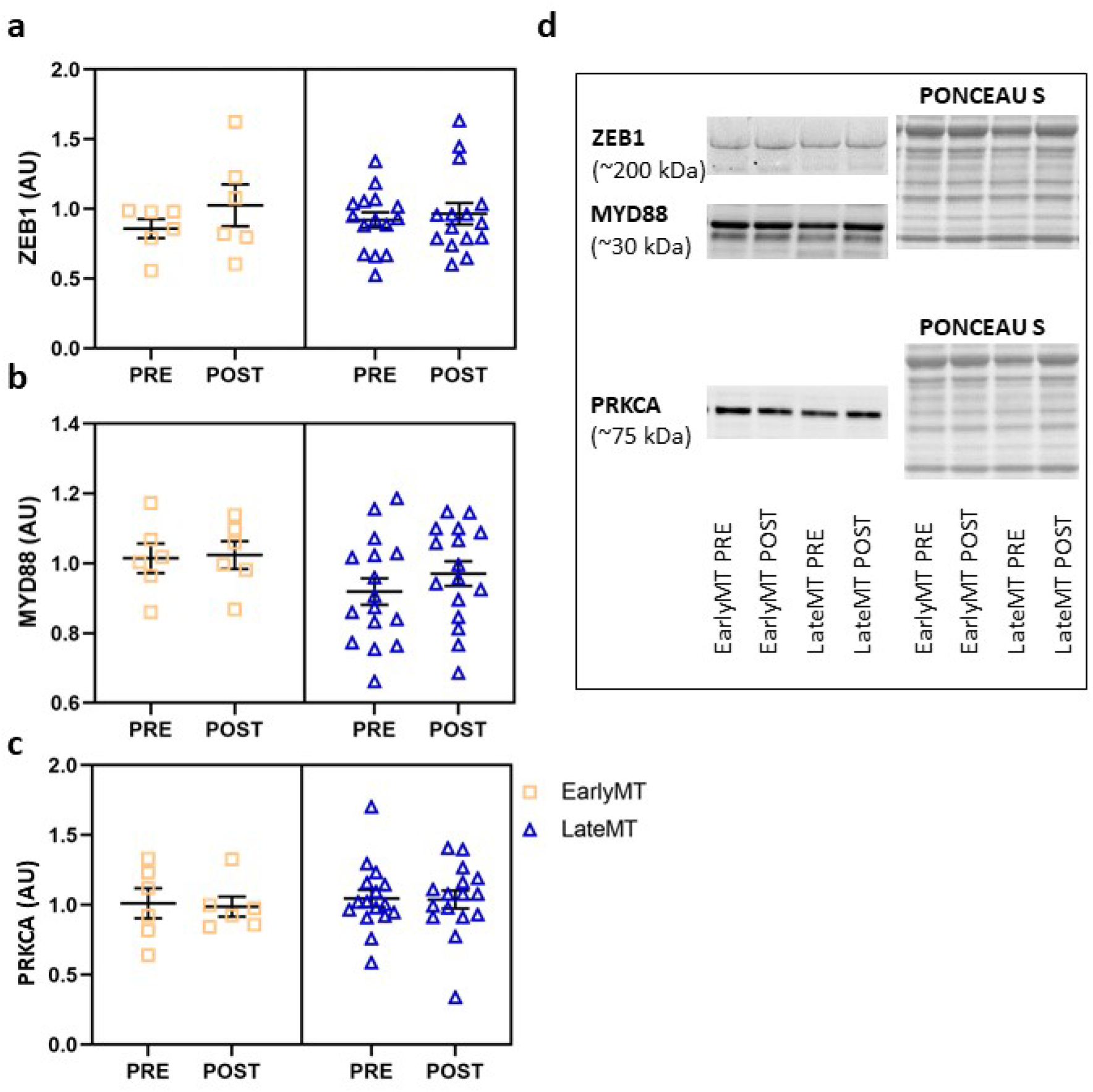
**Western blot results** for **a** ZEB1, **b** MYD88 and **c** PRKCA with **d** representative blot images. Abbreviations: AU, arbitrary units; EarlyMT, a group of women transitioning from early perimenopause to early postmenopause; LateMT, a group of women transitioning from late perimenopause to early postmenopause; PRE, baseline sample; POST, follow-up sample.

### Changes in mRNA gene and regulatory RNA expressions correlated with the changes in body composition variables

Next, we investigated whether the changes (Δ) in muscle RNA expression correlated with the measured changes in body composition (**Figure 5**). The potential “muscularity regulators” in EarlyMT were Δ*GTF2F2* and Δ*C1QTNF9*, which were positively correlated with lean and muscle mass variables, and Δ*TMEM39B*, which was negatively correlated. The potential “adiposity regulators” in EarlyMT were Δ*INTU*, Δ*C1QTNF9* and Δ*PIDD1*, which were positively correlated with adiposity variables, and Δ*ALMS1*, Δ*EXTL3* and Δ*MAFK*, which were negatively correlated. In LateMT, all correlations were rather weak likely because the phenotype-level changes were also modest in the LateMT group.

**Figure 5.**
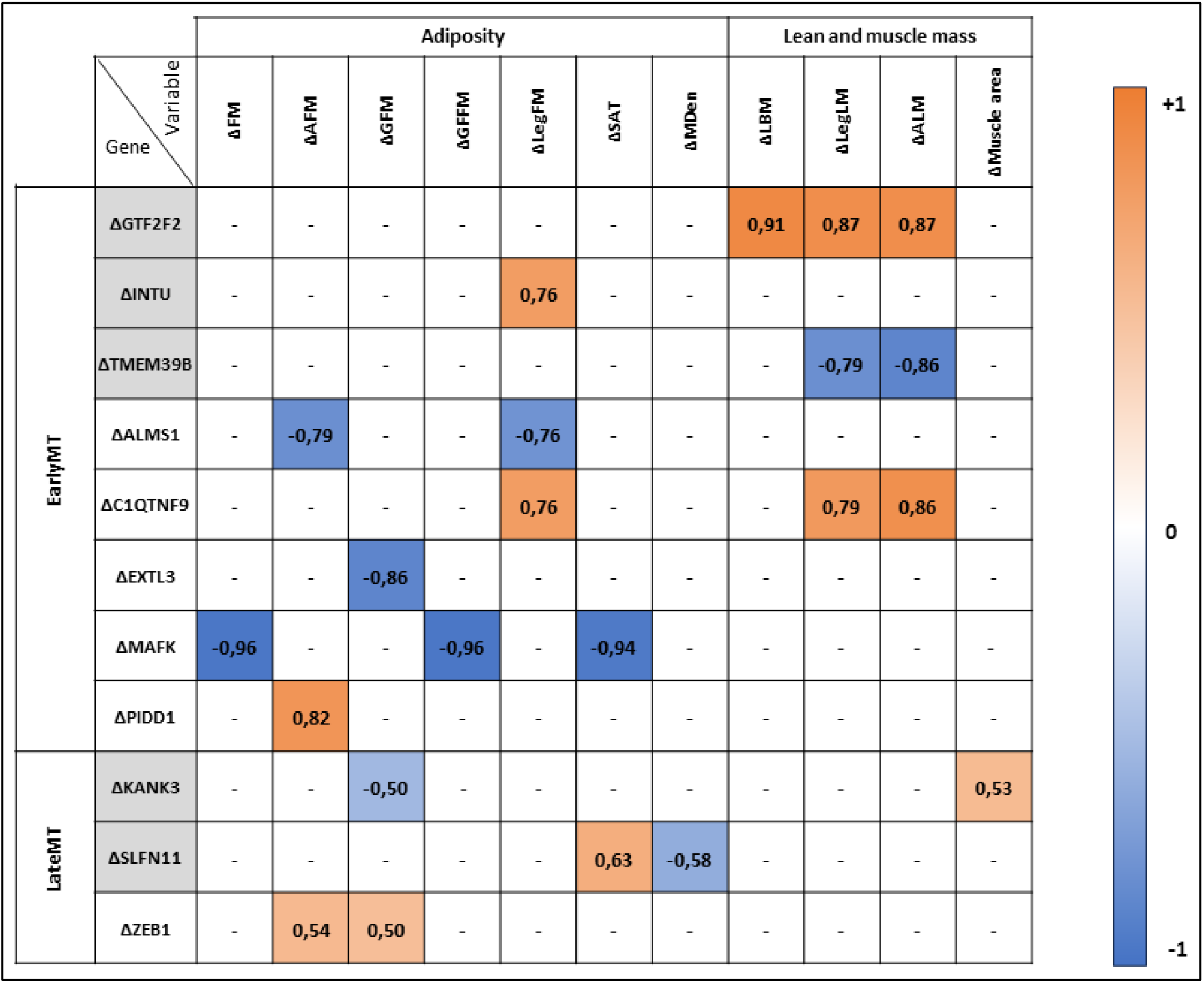
Significant (p < 0.05) correlations of changes in the differentially expressed mRNA genes with the body composition variables. Grey gene, upregulated in postmenopause; white gene, downregulated in postmenopause; orange, positive correlation; blue, negative correlation. FM, total fat mass; AFM, android fat mass; GFM, gynoid fat mass; GFFM, gluteofemoral fat mass; LegFM, right leg fat mass; SAT, cross-sectional subcutaneous adipose tissue area of the thighMDen, muscle density; LBM, total lean body mass; LegLM, right leg lean mass; ALM, appendicular lean mass; Muscle area, cross-sectional muscle area of the thigh.

Although we found no DE regulatory RNAs at gene level, we nevertheless investigated whether the expression level changes in the top 20 expressed lncRNAs genes and miRNAs correlated (r > ± 0.49 and p < 0.05) with body composition variables. In EarlyMT, several positive correlations with adiposity variables were found for ΔMIR1-1HG and negative correlations for ΔAC087190.3, ΔmiR-21-5p, -99a-5p and -126-3p. For lean mass variables, several positive correlations were found for ΔAC093010.2, and ΔmiR-30d-5p, -206 and -378a-3p (**Supplementary Table 10**). In LateMT, positive correlations were found between at least two adiposity variables and ΔAC068700.1, ΔAC069360.1, ΔMALAT1, ΔmiR-1-3p, -26-5p and -30d-5p. Positive correlations with lean and muscle mass variables were found for ΔAC087190.3 and ΔKCNQ1OT1, and negative with ΔSNHG5 (**Supplementary Table 10**).

## Discussion

This study represents the first longitudinal investigation into the molecular pathways behind skeletal muscle changes during menopausal transition. We investigated longitudinal changes in the muscle transcriptome during the menopausal transition in middle-aged women and their relationship with changes in body composition. Leveraging a multi-omics approach, we identified gene expression enrichment in cellular processes controlling cell survival, growth, and cellular interactions with the external environment. Of the expressed mRNA genes, 49 were differentially expressed between peri- and postmenopausal stages.

Additionally, we observed potential regulatory relationships between representatives of the three RNA classes. Furthermore, IPA predicted female hormones to be upstream regulators of some of the DE mRNAs and regulatory RNAs pointing to cell death, cell proliferation, and glucose metabolism to be affected by menopausal transition in skeletal muscle. Lastly, we investigated whether the observed changes in gene expression were linked to the observed changes in body composition and identified putative muscularity regulators, such as *GTF2F2* and *TMEM39B*, and putative adiposity regulators, such as *ALMS1* and *MAFK*.

The DE mRNA genes that we identified, were related to several important functions in the tissue, such as extracellular matrix modification, muscle cell proliferation, energy metabolism and apoptosis signaling. These findings are partially supported by earlier in vivo studies, which have reported that loss of estrogens reduces oxidative capacity and respiratory function, and activates apoptotic signaling (Campbell et al., 2003; Karvinen et al., 2021; La Colla et al., 2017; Torres et al., 2018). More importantly, our results share similarities with the previous cross-sectional studies conducted among postmenopausal women, which have observed responses in genes related for example to cellular and environmental interactions, anatomical structure, protein post-translational modifications, proteolysis, peptidolysis, and cell proliferation (Pöllänen et al., 2007; Ronkainen et al., 2010). However, we found only one common DE gene (*ZNF84*) among our and the earlier reported findings, despite the similarities in the pathways. ZNF84 has been previously associated with the upregulation of cellular senescence (Strzeszewska-Potyrała et al., 2021). Contrary to our observation showing *ZNF84* to be downregulated during the transition from early perimenopause to early postmenopause, Pöllänen et al (Pöllänen et al., 2007), instead, reported an upregulation of *ZNF84* with deepening postmenopausal status. In that study, participating women were early postmenopausal at the time of baseline muscle sampling, and the follow-up samples were taken 12 months later. Therefore, it is likely that the differences between the identified DE genes despite the similarities of the pathways found in our and other human studies, are explained by differences in the menopausal statuses. The earlier studies have focused on postmenopause and the postmenopausal use of hormone therapy, while here we focused on the natural menopausal transition from peri- to postmenopause without the use of any sex hormone-containing medication for contraception or to treat menopausal symptoms.

Our results suggest that hormonal changes related to menopausal transition affect specifically the mRNA transcriptome of skeletal muscle in middle-aged women. We found, for example, *E2F3* and *ZEB1* to be downregulated and *PRKCA*, *MYD88* and *JAK2* to be upregulated during menopausal transition. Supporting the hypothesis that hormonal change might be the driving force behind this observation, earlier studies have shown, and our IPA analysis suggested, them to be responsive for E2 or P4 with the exception that ZEB1 was not identified by IPA. E2F3 is a transcription factor, which controls genes related to cell cycle, proliferation and apoptosis, and has been classified as a transcriptional activator(Leone et al., 1998). In myoblasts and *in vivo*, E2F3 depletion has been associated with reduced cell proliferation, total lean mass and muscle power (Kim et al., 2019; Song et al., 2015). ZEB1 is required for skeletal muscle regeneration owing to its important role in satellite cell quiescence (Siles et al., 2019). *E2F3* and *ZEB1* have previously been shown to be upregulated by E2 and P4 (R. Liu et al., 2011; Mazur et al., 2015; Wu et al., 2018), and their downregulation due to menopause may offer a new pathway for the previously observed menopausal decrease in satellite cells and muscle mass (Collins et al., 2019; Juppi et al., 2020). In muscle, PRKCA has been observed to regulate hypertrophy and to inhibit glucose intake (Braz et al., 2002; Letiges et al., 2002). E2 has been found to upregulate *PRCKA* in breast cancer (Boyan et al., 2003), while our results for muscle suggest the opposite. MYD88, which has a role in myoblast fusion and tissue inflammation, has also been shown to be vital for satellite cells, as its deletion results in loss of muscle mass and strength in rodents (Gallot et al., 2018). MYD88 expression has been shown to be E2 and P4 responsive in monocytes and breast cancer cells (El Sabeh et al., 2021; Matias et al., 2021). Another upregulated gene, JAK2 is a part of the JAK/STAT signaling pathway. In muscle, JAK2 has been associated with the differentiation of satellite cells and improved energy metabolism (J. Liu et al., 2012; K. Wang et al., 2008). E2 has been reported to affect JAK2 activation in a cell type-specific manner (Gupta et al., 2012; Leung et al., 2003).

It is likely that E2 and P4 like other sex hormones exert their effects on gene expression through their receptors. Although not differentially expressed, we observed estrogen receptor ESR1 to be more abundantly expressed than progesterone receptor PGR, thus the loss of E2 may be one of the strongest hormonal contributors to female muscle RNA signaling. In mice, E2 has been shown to regulate the expression of the estrogen receptors (Baltgalvis et al., 2010). Conflicting with these rodent results, but similar to our findings, the earlier human study also reported that systemic E2 levels did not affect muscle estrogen receptor mRNA levels in postmenopausal women (Pöllänen et al., 2007). Why this occurs in rodents, but not in humans, merits more attention. Of the observed 49 DE genes, none were differentially expressed in both EarlyMT and LateMT groups. This further supports the notion that the changes in the skeletal muscle transcriptome in menopause occur in a staged chronological order, possibly reflecting the magnitude of change in E2, which was reduced by nearly 50% from the baseline in EarlyMT while change was negligible in LateMT, or that some threshold level has been crossed. Overall, we observed more changes in mRNA gene expression during the early perimenopause to early postmenopause transition. The duration of the follow-up period did not vary significantly between the two peri-groups, thus, merely time or chronological aging are not the probable contributing factors.

Although we discovered several DE mRNA genes, we did not observe statistically significant changes in the levels of regulatory RNAs. Previous literature from especially muscle lncRNA expression is limited. To our knowledge, muscle lncRNA regulation by menopausal hormones has not been studied in humans before and the corresponding animal studies are rare. In rats, ovariectomy was found to associate with 13 DE muscle lncRNAs (Chai et al., 2019), whereas, in rainbow trout, E2 exposure was found to induce expression level changes in seven lncRNAs (J. Wang et al., 2017). In human cells outside muscle tissue, E2 has been found to induce, for example, H19 expression and repress MALAT1 expression (Sedano et al., 2020). These lncRNA genes were also included in our top 20 expressed lncRNAs in both EarlyMT and LateMT and in fact, we did observe one transcript of MALAT1 (ENST00000618227) to be downregulated in postmenopause. MALAT1 is known to have a role as a promotor of myoblast proliferation and to be a target of myostatin (Watts et al., 2013). MALAT1 is also a target of miR-133 and this connection has been found to regulate myocyte differentiation(Han et al., 2015). Overall, our results somewhat support the literature-driven hypothesis that ovarian hormones might have a regulatory role in muscle lncRNAs, although we observed only mild menopausal changes in the lncRNA gene expression levels. Plausible reasons for the difference may again include different study designs, but also differences in species-specific gene regulation.

To our surprise, we did not observe statistically significant menopausal transition-associated changes in the skeletal muscle miRNA expression. This contrast earlier studies in which we and others have shown E2 levels to influence skeletal muscle miRNA expression. For example, in a study with postmenopausal twins discordant for HT, we showed that the use of HT was associated with lower expression levels of miRNAs miR-142-3p,-182 and -223 in muscle (Olivieri et al., 2014). These miRNAs regulate, for example, the FOXO and IGF1 pathways controlling muscle atrophy and insulin sensitivity. In animal studies, lower E2 levels have been associated with lower expression levels of apoptosis-linked miR-122-5p and -214-3p, and cell proliferation- and differentiation-related miR-26b, -27a-5p, -27b and -199a-3p levels (Karvinen et al., 2021; Martignani et al., 2019). In addition, other studies with *in vitro* models have found E2 and P4 to regulate the expression of several other miRNAs (Bhat-Nakshatri et al., 2009; Klinge et al., 2010; Pan et al., 2017). Of the mentioned miRNAs, in EarlyMT, the expression of miR-26b, -27b, -122-5p, -142-3p and -199a-3p was lower in postmenopause than in perimenopause, i.e., directionally same as in previous studies, and that of miR-27a-5p and -214-3p higher, i.e., contradicting expression concordance as in previous studies, although in our hands none of the differences was statistically significant. Thus, our results are somewhat unexpected when compared to previous studies, although we are aware of the uniqueness of the current dataset. One reason for the discrepancy in results may be that the women in the current study were closer to each other in their hormone levels, especially in E2 levels and the E2 is the body’s own endogenous product rather than an exogenous medical product.

In an attempt to reveal new potential contributors to muscle mass and adiposity, and thereby, metabolic health, we inspected if the change in DE mRNA genes correlated with the observed change in muscularity or adiposity variables. Correlations with changes in several lean and muscle mass variables were found for Δ*GTF2F2*, Δ*C1QTNF9* and Δ*TMEM39B*. GTF2F2 encodes a polypeptide that is a component of the transcription-initiating complex but has not been reported to be functional in skeletal muscle. In skeletal muscle, loss of C1QTNF9 protein leads to reduced insulin signaling and mitochondrial content (Wei et al., 2014). In cardiac myocytes and smooth muscle, it reduces apoptosis (Kambara et al., 2012), but also cell proliferation (Uemura et al., 2013). Thus, although data from human skeletal muscle are not yet available, the results on other cell types suggest, that the downregulation of this protein may contribute to cell loss.

This notion is supported by our correlation analyses, which showed a positive association between the change in *C1QTNF9* level and the changes in lean mass variables. TMEM39B is a transmembrane protein, which regulates the production and secretion of procollagen and regulates endoplasmic reticulum stress (Zhang et al., 2021), but is still poorly understood in skeletal muscle. Interestingly, in a study of older men and women, *TMEM39B* was found to be upregulated in older age (Thalacker-Mercer et al., 2010), as in our data. In our study, the change in the expression of *TMEM39B* was also negatively associated with change in leg (ΔLegLM) and appendicular lean mass (ΔALM), indicating that it may have a function in the loss of muscle mass, although this may be related also to aging. Among the adiposity variables, we found significant negative correlations for Δ*ALMS1* and Δ*MAFK.* ALMS1 has a function in cell microtubule organization and defects in *ALMS1* have been associated with young-onset obesity syndrome (Collin et al., 2002). In epithelial cells, E2 decreases *ALMS1* mRNA expression (Winuthayanon et al., 2014), but no studies thus far have reported its role in skeletal muscle. In our study, *ALMS1* was downregulated in postmenopause, and the change correlated negatively with change in android and leg fat mass. This might imply that in skeletal muscle, the loss of estrogen signaling decreases *ALMS1* expression, which then further contributes to increased adiposity. In addition, ALMS1 expression was also found to correlate directly with the expression of positive contributors to muscle mass myomiRs miR-486- 3p and -5p. *MAFK*, a transcriptional repressor, has been found to be upregulated by progestogens in endometrial stromal cells (Vallejo et al., 2014), but information from muscle is lacking. We found *MAFK* to be downregulated in postmenopause and to have very strong negative associations with body adiposity variables. How these two observations are connected remains to be investigated.

Overall, we conducted a thorough investigation of the effects of the menopausal transition on the skeletal muscle transcriptome. To our knowledge, this is the first study to investigate these associations. However, the study has its limitations. One limitation is the relatively small sample size. Moreover, this study focused on RNA expression and was only able to investigate the protein-level effects of three genes. The strengths of the study include the longitudinal study design and the use of gold-standard methods in menopausal status assignment and body composition measurements.

## Conclusions

The menopausal transition was associated with changes in human skeletal muscle transcriptome specifically regarding mRNA gene expression. The non-protein coding regulatory RNAs, i.e., lncRNA genes or miRNAs, showed only subtle changes in their expression or higher variation between samples prohibiting statistical significance from being observed. Nevertheless, suggestive regulatory networks were identified. The DE mRNA genes identified in this study may contribute to muscle tissue homeostasis and further the unfavorable menopausal changes in total body composition. Moreover, the observed DE mRNA genes differed between the two perimenopausal groups, which may indicate change in the regulatory mechanisms at different stages of the menopausal transition. The observed DE mRNA genes serve as a possible starting point for mechanistic studies aimed at a more detailed understanding of the effects of hormonal changes in female skeletal muscle.

## Material and methods

### Study design and population

This study used the data from the ERMA (Estrogenic Regulation of Muscle Apoptosis) study, in which originally 6878 women aged 47–55 years were approached with questionnaires in 2014 (Kovanen et al., 2018; Laakkonen et al., 2022). Of them, 1393 women provided blood samples and menstrual cycle diaries and were assigned to four menopausal groups (premenopausal, early perimenopausal, late perimenopausal and postmenopausal) based on modified STRAW+ 10 guidelines (Harlow et al., 2012). A subgroup of perimenopausal women (n = 381) consented to take part in the ERMA longitudinal study, in which they were monitored over the menopausal transition to early postmenopause. As the process of menopausal transition varies from one woman to another, the follow-up time was individualized accordingly.

At all laboratory visits, fasting serum samples were taken from the antecubital vein between 7–10 am. Progression of the menopausal transition was monitored by assessing serum FSH levels every three to six months and only after two elevated FSH measurements (>30 IU/L) accompanied by at least six months of self-reported lack of menstruation was a participant considered to be early postmenopausal. The participant was then invited to the final follow-up visit for physiological measurements during which her E2 and FSH levels were again measured. To minimize the effect of daily fluctuation in hormone levels, we use an average of the two last measurements as the follow-up E2 and FSH value. Serum E2 and FSH were measured with IMMULITE 2000 XPi (Siemens Healthcare Diagnostics, UK).

Muscle biopsies of 25 women (EarlyMT, n = 8 and LateMT, n = 17) were collected at baseline and final follow-up measurements. None of the women used estrogen- or progestogen-containing hormonal therapy during the study. Muscle biopsies were collected from the middle part of *m. vastus lateralis* using the Bergström needle biopsy method under local anesthesia. All visible connective and adipose tissue were removed, and the sample was snap-frozen in liquid nitrogen inside aluminum foil wrap. Samples were stored at -150°C until analysis. The samples of one EarlyMT participant were later excluded from analysis due to low RNA/sequencing data quality (see *RNA isolation and sequencing*).

The study was performed in accordance with the Declaration of Helsinki. All participants provided their written informed consent, and the study was approved by the ethical committee of the Central Finland Health Care District (ERMA 8U/2014).

### Anthropometrics and body composition

Body mass was measured with a digital scale and BMI was calculated by dividing body mass (kg) by height (m) squared. Body composition was measured with dual-energy X-ray absorptiometry (DXA, LUNAR Prodigy, GE Healthcare, Chicago, IL, USA). Measures of total (LBM), appendicular (ALM) and right leg lean mass (LegLM) and total fat mass (FM), total fat percentage (Fat%), android (AFM), gynoid (GFM), gluteofemoral (GF-FM) and right leg fat mass (LegFM) were analyzed from the scans. The gluteofemoral area was outlined manually using the iliac crest line as the upper limit and the mid-knee joint as the lower limit of a rectangle as reported previously (Juppi et al., 2022). All measurements were done in light underwear and after overnight fasting.

In addition to DXA, the right mid-thigh was scanned at the level of the muscle biopsy with quantitative computed tomography (qCT, Siemens Somatom Emotion scanner, Siemens, Erlangen, Germany). The areas of subcutaneous adipose tissue (SAT area) and muscle (Muscle area) of the thigh were measured using appropriate thresholds in Python Software (version 3.6). From the cross-sectional image, the muscle portion including the femur was first separated using a U-net machine learning algorithm or, if needed, manually.

Adipose tissue, muscle and bone area were separated using Hounsfield Unit (HU) limits and the mean density of muscle tissue (MDen) was calculated. Images were analyzed using ImageJ Software (v.1.52, NIH, USA) and Python.

### Background information

Information about smoking (current/non-smoker), alcohol consumption (units per week), medical conditions, prescription medication and the use of products containing exogenous sex hormones was determined from questionnaires. Preparations containing estrogen alone or combined with progestogen including transdermal (patches, gels and sprays), oral (tablets) and intrauterine hormonal preparations were regarded as hormone therapy, but local intravaginal estrogen therapy was not. Diet was assessed using food-frequency questionnaires and quantified using the diet quality score (DQS) as reported previously (Juppi et al., 2020). The amount of physical activity was assessed from the structured questionnaire and with hip-worn accelerometers (Hyvärinen et al., 2021).

### RNA isolation and sequencing

Total RNA from ∼10–60mg of muscle tissue was extracted using a Qiagen miRNeasy Mini kit (217004, Qiagen, Hilden, Germany) according to the manufacturer’s instructions. In brief, tissue was weighted and homogenized in QIAzol Lysis Reagent for two minutes with 25hz, using TissueLyser and a sterile metal bead. Total RNA was extracted according to the manual and eluted to 30µl of RNase-free water. Samples were immediately stored at -80°C until analysis.

*Sequencing* was outsourced to the Novogene (Novogene Company Limited, Cambridge, United Kingdom). For the long RNA transcripts (mRNA and lncRNA), library construction and quality control were performed according to the Novogene protocols. In brief, ribosomal RNA was removed from the samples and the remaining RNA was fragmented. Single complementary DNA strands were then synthesized with reverse transcription. A mixture of dNTPs, RNAse H and DNA polymerase I was added to initiate second-strand synthesis. After a series of end repairs, A-tailing and the use of a U-adaptor, PCR amplification resulting in a final double-stranded cDNA library was performed. Libraries were analyzed with agarose gel and correct-size libraries were selected for sequencing. After pooling, sequencing was performed using Illumina NovaSeq 6000 at Novogene and paired-end reads of 150bp were generated. The average sequencing depth was 35M mapped reads per sample.

For the small RNA (sRNA) library, sequencing libraries were generated using a NEBNext® Multiplex Small RNA Library Prep Set for Illumina® (NEB, USA) following the manufacturer’s recommendations. Index codes were added to attribute sequences to each sample. PCR amplification was performed using LongAmp Taq 2X Master Mix, SR Primer for Illumina and index (X) primer. PCR products were purified on polyacrylamide gel and DNA fragments corresponding to ∼140–160 bp (the length of small noncoding RNA plus the 3’ and 5’ adaptors) were recovered and dissolved in the elution buffer. Finally, library quality was assessed on the Agilent Bioanalyzer 2100 system using DNA High Sensitivity Chips. RNA sequencing was performed using Illumina NovaSeq 6000 at Novogene and 50bp single-end reads were generated. The average sequencing depth was 10.2 mapped reads per sample.

*Quality control.* Before sequencing, RNA integrity, quantitation and sample purity were assessed using the RNA Nano 6000 Assay Kit of the Agilent Bioanalyzer 2100 system (Agilent Technologies, CA, USA) and agarose gel electrophoresis. The integrity value of all samples was between 7 and 9.1, the mean integrity value being 8.6. After sequencing, the mean Q30 value was on average 96% for sRNA and 94% for mRNA and lncRNA, indicating high-quality data. The sequence length distribution of sRNA was then checked for each sample to ensure enrichment of miRNAs. Due to distorted sequence length distribution and low RNA integrity value compared to other samples, one EarlyMT sample pair was excluded from all sequence data analysis. Thus, the final N of samples selected for further analysis was 48, comprising 24 longitudinal sample pairs (for EarlyMT n = 7 sample pairs and for LateMT n = 17 sample pairs).

### Data analysis

*Preprocessing of long RNAs.* For long transcripts, raw data were first processed through Novogene in-house scripts to identify mRNAs and lncRNAs. Reads containing adapter, poly-N sequences or with low quality were removed. Q20, Q30 and GC content of the clean data were calculated. Clean paired-end reads were mapped to the reference genome (Hg38) using HISAT2 software. The mapped reads of each sample were assembled into potential transcripts by StringTie and merged through Cuffmerge. To identify novel lncRNAs transcripts (TCONS_), the transcripts were analyzed for evidence of their protein-coding potential and their conservation of sequence with known proteins. To identify lncRNAs, low expression level transcripts were filtered out, the exon number was set to >2 and the transcript length was set to >200nt. CNCI (Coding-Non-Coding-Index), CPC (Coding Potential Calculator) and Pfam were used to predict the coding potential of the transcripts. If at least one of the three tools predicted a transcript to have coding potential, it was filtered out and considered to represent mRNA transcript. The transcripts that were not identified to have coding potential with any of the three prediction tools formed the candidate transcript set of lncRNAs. Finally, transcripts were compared and evaluated against a reference genome using Cuffcompare. A filter was applied before DE analysis of mRNA and lncRNA. *mRNA-filter:* To be included in the gene-level analysis, the mRNA transcript had to be expressed a minimum of one count per million (CPM>1) and to be present in 21% of the samples of both groups (3/14 samples in EarlyMT or 7/34 samples in LateMT). In addition, based on Ensembl database transcript biotype, we excluded transcripts without a valid Ensembl ID and no reference for protein-coding ability. *lncRNA-filter:* equivalent to mRNA-filter, lncRNA transcript needed to be expressed at level CPM>1 and present in 21% of the samples. The transcript also had to have valid ENST- and ENSG-IDs and the Ensembl biotype of “lncRNA”. For both mRNA and lncRNA, to correct for potential differential isoform usage, *tximport* R package was used to transform the transcript-level data into a format that could be input to *DESeq2*. *Tximport* calculates gene-level estimated counts and a gene-level offset that corrects for differences in the average transcript length across samples (Soneson et al., 2016).

*Preprosessing of short RNAs.* The raw sRNA sequence data was first processed through Novogene in-house scripts. The data was cleaned by removing reads with poly-N, 5’ adapter contaminants, without 3’ adapter or the insert tag, reads containing poly-A/-T/-G or C, and low-quality reads. The sRNA (18–35bp) sequences were mapped to the reference sequence (Hg38) using Bowtie. miRBase20.0 was used as reference, modified software mirdeep2 and srna-tools-cli were used to obtain the potential miRNA. Custom scripts by Novogene were used to obtain the miRNA counts. Equivalent to mRNA and lncRNA DE analysis, a filter was applied before miRNA DE analysis. *miRNA-filter:* The included miRNAs had to have CPM>1 and to be present in at least 21% of the samples per group.

*The DE analysis of long and short RNAs.* DE analysis was performed with R package *DESeq2*(Love et al., 2014) with paired-samples design formula and default settings. Genes, transcripts or miRNAs with LFC > ± 1.5 and p_adj_< 0.05 (p_adj_ = p-value subjected to FDR correction) were regarded as differentially expressed.

### Gene Enrichment analysis

Unique gene enrichment in the EarlyMT (n = 314 mRNA genes) and LateMTa (n = 175 mRNA genes) groups were investigated using Panther 18.0 (pantherdb.org) against GO Biological Processes databases. The main transcripts (highest expression level) of all the prefiltered mRNA genes (for EarlyMT n = 12,328 and LateMT n = 12,189) were imported into GSEA (Gene Set Enrichment analysis). Analysis was performed with the *fgsea*(Korotkevich et al., 2016) R package using t-test statistic (difference in means scaled by the standard deviation and number of samples) as the ranking metric. Reactome and GO Biological Processes databases from Molecular Signatures Database (MSigDB) were used as data sources.

### RNA network interaction

Associations between the different RNA species were investigated with *mixOmics* (Rohart et al., 2017) R package using multiblock sPLS-DA. Datasets were constructed from normalized EarlyMT and LateMT expression data by dividing the postmenopause values by the perimenopause values for the same individual. mRNA input data was obtained from the DE genes. For the lncRNAs and miRNAs, the input data was obtained from the 20 most abundantly expressed, 20 most absolutely and 20 most relatively differentially expressed lncRNA transcripts or miRNAs in each menopausal group. EarlyMT and LateMT datasets were compared to each other with block sPLS-DA. The number of variables allowed to be selected for the analysis were 10 miRNAs, 10 mRNAs and 10 lncRNAs. The resulting construction of the relevance network was plotted using correlation cutoff value 0.7 and re-plotted with Cytoscape.

Interactions between upstream regulators, mRNAs, lncRNAs, miRNAs and downstream functions were investigated using IPA (Qiagen). For the core analysis of IPA, all RNA molecules with unadjusted p < 0.05 and LFC ≥ ± 1.5 were included (for EarlyMT n = 267 and n = 133 for LateMT). For the My pathway analysis of IPA, we selected the DE mRNA genes (**Table 3**) and the top 20 abundantly expressed (normalized base mean per group) lncRNA genes and miRNAs.

### Western blot

Total protein from muscle tissue samples (n = 6 for EarlyMT and n = 16 for LateMT) was extracted using an extraction buffer containing Tissue extraction Reagent I (FNN0071, Invitrogen, Waltham, Massachusetts, USA), HALT Protease and Phosphatase Inhibitor Cocktail (1:100, 1861280, Thermo Scientific, Rockford, IL, USA), pepstatin A (1:100, P5318, Sigma-Aldrich, Saint Louis, Missouri, USA) and 0.5M EDTA (1:100). Frozen tissue was lysed in Tissuelyzer 2x2 min 25 Hz or until the solution was homogenous. Homogenates were first shaken for 30 minutes and then centrifuged for 10 minutes at 10000xg at +4°C to gain protein-containing supernatant. Protein concentration was measured using Pierce BCA Protein assay kit (23225, Thermo Scientific) in Indiko Plus (Thermo Scientific). 25ug of protein was used for Western blot run in 4-20% gradient gel (Criterion TGX Stain-free, #5678094, Bio-Rad, Hercules, California, USA). Proteins were transferred to nitrocellulose membrane in Turboblotter (Bio-Rad, USA) and total protein concentration of the lanes was measured using standard Ponceau S protocol. Destained membranes were blocked for 2 hours RT in Intercept Blocking Buffer (#927-70001, LI-COR, Lincoln, Nebraska, USA). Primary antibodies for MYD88 (ab107585, Abcam, Cambridge, UK), ZEB1 (ab203829, Abcam), PRKCA (ab32375, Abcam), JAK2 (ab108596, Abcam), E2F3 (ab152126, abcam) and APC (ab40778, Abcam) were tested for optimal dilutions. Antibodies for JAK2 (tested dilutions 1:2500 and 1:500), E2F3 (tested dilutions 1:1000 and 1:500) and APC (tested dilutions 1:2500 and 1:500) were unspecific or unfunctional. Primary antibodies for MYD88 (1:750), ZEB1 (1:500), and PRKCA (1:5000) were incubated in 1:1 TBS-Blocking buffer-solution in +4°C O/N. The next day, membranes were washed and IRDye secondaries (#926-32213, 800CW Donkey anti-rabbit and #926-32212, 800CW Donkey anti-mouse, both in 1:10000, LI-COR, USA) in 1:1 TBST-Blocking buffer-solution were incubated for 1h RT. Membranes were washed and imaged with ChemiDoc MP (Bio-Rad, USA). Band intensities were analysed in ImageLab (version 6.0.1, Bio-Rad, USA) and first normalized to blot average followed by normalization to the corresponding Ponceau S total protein concentration. Results are expressed in arbitrary units (AU).

### Statistical analysis

All variables were evaluated for normality and parametric tests were used where appropriate. Longitudinal changes in characteristics were investigated using paired T-test and Wilcoxon Signed Rank test. Differences in changes between EarlyMT and LateMT groups were investigated using independent samples T-test and Mann-Whitney U-test. Western blot results were investigated using the Wilcoxon Signed Rank Test. The correlations between the changes in gene/transcript expression level and body composition variables (Δ = POST minus PERI) were calculated using Spearman correlations. For mRNAs and lncRNA genes, the most abundantly expressed transcript was used in correlation analysis.

## Data availability

The sequencing data is available from the NCBI Sequence Read Archive (SRA, data-ID will be updated here) and the main code used to conduct this study is available on GitHub at <u>LaakkonenLab/menopausal-</u> <u>muscle-RNA (github.com).</u> Other datasets including the source data supporting our findings are shared as supplementary files with this paper. The complete individual-level data is not publicly available due to privacy or ethical restrictions. Access to the data may be granted upon justification for research purposes taken that a request does not violate the consent provided by the study participants. Such requests should be addressed to the project leader and will be evaluated with the board consisting of senior researchers involved in the ERMA study. The metadata of the ERMA study is publicly available (doi:10.17011/jyx/dataset/83491).

## Acknowledgements

This study was funded by grants from the Academy of Finland (grant numbers 275323 to V.K. and 309504, 314181, and 335249 to E.K.L). We thank Mervi Matero, Hanne Tähti and all other laboratory staff at the Faculty of Sport and Health Sciences for their invaluable assistance with data collection. We also thank the women who participated in the ERMA study for donating their time and effort.

## Author contributions

V.K., E.K.L., and S.S. designed the original ERMA study and U.M.K., and P.A. contributed to the planning. E.K.L. and H-K.J. were responsible for the muscle sample handling. H-K.J. did RNA and protein isolations and related work while S.K. performed the Western Blot and related analyses. H-K.J. and N.C. analysed the DXA and CT scans. S.S. performed the CT scanning and supervised the body composition analyses. H-K.J did body composition-related correlation analysis. H-K.J did the majority of the pre-analysis of datafiles provided by Novogene, performed IPA and constructed the final tables while T-M.K. performed the main bioinformatic analyses and result images, while T.S. assisted. V.K. and E.K.L. provided funding for the study. H-K.J. prepared the first version of the manuscript. H-K.J., T-M.K., T.S., V.K., U.M.K., P.A., N.C., S.S., S.K., and E.K.L have participated in the interpretation of the results and critically commented on the manuscript during the writing process. E.K.L, H-K.J and T-M.K prepared the final version of the manuscript, which all the authors have read and approved.

## Material and correspondence

Eija K. Laakkonen

## Ethics and participant consent

The study was performed in accordance with the Declaration of Helsinki. All participants provided written informed consent, and the study was approved by the ethical committee of the Central Finland Health Care District (ERMA 8U/2014 and EsmiRs 9U/2018).

## Permission to reproduce material from other sources

Not applicable.

## ABBREVIATIONS USED FOR GENES AND PROTEINS

ACTA1: actin
ALMS1: ALMS1 centrosome and basal body associated protein APC, adenomatosis polyposis coli
AR: androgen receptor
BAIAP2-DT: BAR/IMD domain containing adaptor protein 2 divergent transcript BIRC6, baculoviral IAP repeat containing 6
C1QTNF9/ CTRP9: C1q And TNF related 9
CTRP15/myonectin: Complement C1q Tumor Necrosis Factor-Related Protein 15 CYP19A1, aromatase
E2F3: E2F transcription factor 3
ECM1: extracellular matrix protein 1 ELN, elastin
ENTPD1-AS1: ectonucleoside triphosphate diphosphohydrolase 1 antisense RNA 1 ESR1/ERα, estrogen receptor 1
ESR2/ERβ: estrogen receptor 2
EXTL3: exostosin-like glycosyltransferase 3
FBXO32/Atrogin-: F-Box Protein 32, MAFbx, 1 FNDC5, fibronectin type III domain-containing 5, (irisin) FOXO1, forkhead box O1
FOXO3: forkhead box O3
FSHR: follicle-stimulating hormone receptor
GLUT4/SLC2A4: glucose transporter type 4, solute carrier family 2 (facilitated glucose transporter), member 4
GPER1: G-protein-coupled estrogen receptor 1 GTF2F2, general transcription factor IIF subunit 2 HSD17B1, hydroxysteroid 17-beta dehydrogenase 1 HSD17B2, hydroxysteroid 17-beta dehydrogenase 2 HSD17B4, hydroxysteroid 17-beta dehydrogenase 4 HSD17B5, hydroxysteroid-17-beta dehydrogenase 5 HSD3B2, progesterone reductase
IGF1: insulin-like growth factor 1 IL-15, interleukin 15
INTU: inturned planar cell polarity protein
IQCH-AS1: IQ motif containing H antisense RNA 1 JAK2, janus kinase 2
KCNQ1OT1: potassium voltage-gated channel subfamily Q member 1 opposite transcript 1 KIAA0355/GARRE, granule associated rac and RHOG effector 1
LINC00667: long intergenic non-protein coding RNA 667 LINC02541, long intergenic non-protein coding RNA 2541 MAFK, MAF bZIP transcription factor K
MALAT1: metastasis-associated lung adenocarcinoma transcript 1 MAPK, mitogen-activated protein kinase 1
MGLL: monoglyceride lipase MIR1-1HG, MIR1-1 host gene
MKNK1: MAPK interacting serine/threonine kinase 1 mTOR, mechanistic target of rapamycin kinase
MuRF-1/TRIM63: muscle-specific RING finger protein 1 MYD88, myeloid differentiation primary response protein MyD88 MYH1, myosin heavy chain 1 (IIX)
MYH2: myosin heavy chain 2 (IIA) MYH7, myosin heavy chain 7 (I) MyoD, myogenic differentiation 1
NAA35: N-alpha-acetyltransferase 35, NatC auxiliary subunit NEAT1, nuclear-enriched abundant transcript 1
NEB: nebulin
NORAD: non-coding RNA activated by DNA damage NUDT4, nudix hydrolase 4
NUTM2A-AS1: NUT family member 2a antisense RNA 1
OSER1-DT: oxidative stress responsive serine rich 1 divergent transcript PGR, progesterone receptor
PI3K/Akt: phosphatidylinositol 3-kinase/Akt serine/threonine kinase PIDD1, p53-induced death domain protein 1
PRKCA/PKCa: protein kinase C alpha
RBMS1: RNA binding motif single-stranded-interacting protein 1 SGSH, N-sulfoglucosamine sulfohydrolase
SNHG14: small nucleolar RNA host gene 14 SNHG5, small nucleolar RNA host gene 5 SRD5A1, steroid 5-alpha-reductase 1 TMEM39B, transmembrane protein 39B TTN, titin
XIST: X inactive specific transcript
ZEB1: zinc finger E-box-binding homeobox 1 ZNF611, zinc finger 611
ZNF710-AS1: zinc finger 710 antisense RNA 1 ZNF84, zinc finger 84

## References

Abkhooie, L., Sarabi, M. M., Kahroba, H., Eyvazi, S., Montazersaheb, S., Tarhriz, V., & Hejazi, M. S. (2021). Potential Roles of MyomiRs in Cardiac Development and Related Diseases. Current Cardiology Reviews, 17(4), e010621188335. 10.2174/1573403X16999201124201021

Baltgalvis, K. A., Greising, S. M., Warren, G. L., & Lowe, D. A. (2010). Estrogen Regulates Estrogen Receptors and Antioxidant Gene Expression in Mouse Skeletal Muscle. PLoS ONE, 5(4), e10164. 10.1371/journal.pone.0010164

Bentzinger, C. F., Wang, Y. X., & Rudnicki, M. A. (2012). Building Muscle: Molecular Regulation of Myogenesis. Cold Spring Harbor Perspectives in Biology, 4(2), a008342. 10.1101/cshperspect.a008342

Bhat-Nakshatri, P., Wang, G., Collins, N. R., Thomson, M. J., Geistlinger, T. R., Carroll, J. S., Brown, M., Hammond, S., Srour, E. F., Liu, Y., & Nakshatri, H. (2009). Estradiol-regulated microRNAs control estradiol response in breast cancer cells. Nucleic Acids Research, 37(14), 4850–4861. 10.1093/nar/gkp500

Bodine, S. C., Stitt, T. N., Gonzalez, M., Kline, W. O., Stover, G. L., Bauerlein, R., Zlotchenko, E., Scrimgeour, A., Lawrence, J. C., Glass, D. J., & Yancopoulos, G. D. (2001). Akt/mTOR pathway is a crucial regulator of skeletal muscle hypertrophy and can prevent muscle atrophy in vivo. Nature Cell Biology, 3(11), Article 11. 10.1038/ncb1101-1014

Bondarev, D., Laakkonen, E., Finni, T., Kokko, K., Kujala, U., Aukee, P., Kovanen, V., & Sipilä, S. (2018). Physical performance in relation to menopause status and physical activity. Menopause (New York, N.Y.), 25(12), 1432–1441. 10.1097/GME.0000000000001137

Boyan, B. D., Sylvia, V. L., Frambach, T., Lohmann, C. H., Dietl, J., Dean, D. D., & Schwartz, Z. (2003). Estrogen-Dependent Rapid Activation of Protein Kinase C in Estrogen Receptor-Positive MCF-7 Breast Cancer Cells and Estrogen Receptor-Negative HCC38 Cells Is Membrane-Mediated and Inhibited by Tamoxifen. Endocrinology, 144(5), 1812–1824. 10.1210/en.2002-221018

Braz, J. C., Bueno, O. F., De Windt, L. J., & Molkentin, J. D. (2002). PKCα regulates the hypertrophic growth of cardiomyocytes through extracellular signal–regulated kinase1/2 (ERK1/2). The Journal of Cell Biology, 156(5), 905–919. 10.1083/jcb.200108062

Campbell, S. E., Mehan, K. A., Tunstall, R. J., Febbraio, M. A., & Cameron-Smith, D. (2003). 17beta-estradiol upregulates the expression of peroxisome proliferator-activated receptor alpha and lipid oxidative genes in skeletal muscle. Journal of Molecular Endocrinology, 31(1), 37–45. 10.1677/jme.0.0310037

Chai, S., Wan, L., Wang, J.-L., Huang, J.-C., & Huang, H.-X. (2019). Systematic analysis of long non-coding RNA and mRNA profiling using RNA sequencing in the femur and muscle of ovariectomized rats. Journal of Musculoskeletal and Neuronal Interactions, 19(4), 422–434.

Collin, G. B., Marshall, J. D., Ikeda, A., So, W. V., Russell-Eggitt, I., Maffei, P., Beck, S., Boerkoel, C. F., Sicolo, N., Martin, M., Nishina, P. M., & Naggert, J. K. (2002). Mutations in ALMS1 cause obesity, type 2 diabetes and neurosensory degeneration in Alström syndrome. Nature Genetics, 31(1), 74–78. 10.1038/ng867

Collins, B. C., Arpke, R. W., Larson, A. A., Baumann, C. W., Xie, N., Cabelka, C. A., Nash, N. L., Juppi, H.-K., Laakkonen, E. K., Sipilä, S., Kovanen, V., Spangenburg, E. E., Kyba, M., & Lowe, D. A. (2019). Estrogen Regulates the Satellite Cell Compartment in Females. Cell Reports, 28(2), 368–381.e6. 10.1016/j.celrep.2019.06.025

El Sabeh, R., Bonnet, M., Le Corf, K., Lang, K., Kfoury, A., Badran, B., Hussein, N., Virard, F., Treilleux, I., Le Romancer, M., Lebecque, S., Manie, S., Coste, I., & Renno, T. (2021). A Gender-Dependent Molecular Switch of Inflammation via MyD88/Estrogen Receptor-Alpha Interaction. Journal of Inflammation Research, 14, 2149–2156. 10.2147/JIR.S306805

Fernandes, J. C. R., Acuña, S. M., Aoki, J. I., Floeter-Winter, L. M., & Muxel, S. M. (2019). Long Non-Coding RNAs in the Regulation of Gene Expression: Physiology and Disease. Non-Coding RNA, 5(1), 17. 10.3390/ncrna5010017

Friedman, R. C., Farh, K. K.-H., Burge, C. B., & Bartel, D. P. (2009). Most mammalian mRNAs are conserved targets of microRNAs. Genome Research, 19(1), 92–105. 10.1101/gr.082701.108

Frontera, W. R., & Ochala, J. (2015). Skeletal muscle: A brief review of structure and function. Calcified Tissue International, 96(3), 183–195. 10.1007/s00223-014-9915-y

Gallot, Y. S., Straughn, A. R., Bohnert, K. R., Xiong, G., Hindi, S. M., & Kumar, A. (2018). MyD88 is required for satellite cell-mediated myofiber regeneration in dystrophin-deficient mdx mice. Human Molecular Genetics, 27(19), 3449–3463. 10.1093/hmg/ddy258

Greendale, G. A., Sternfeld, B., Huang, M., Han, W., Karvonen-Gutierrez, C., Ruppert, K., Cauley, J. A., Finkelstein, J. S., Jiang, S.-F., & Karlamangla, A. S. (2019). Changes in body composition and weight during the menopause transition. JCI Insight, 4(5), e124865. 10.1172/jci.insight.124865

Gupta, N., Grebhardt, S., & Mayer, D. (2012). Janus kinase 2—A novel negative regulator of estrogen receptor α function. Cellular Signalling, 24(1), 151–161. 10.1016/j.cellsig.2011.08.016

Han, X., Yang, F., Cao, H., & Liang, Z. (2015). Malat1 regulates serum response factor through miR-133 as a competing endogenous RNA in myogenesis. The FASEB Journal, 29(7), 3054–3064. 10.1096/fj.14-259952

Harlow, S. D., Gass, M., Hall, J. E., Lobo, R., Maki, P., Rebar, R. W., Sherman, S., Sluss, P. M., de Villiers, T. J., & for the STRAW 10, C. G. (2012). Executive summary of the Stages of Reproductive Aging Workshop + 10: Addressing the unfinished agenda of staging reproductive aging. Menopause, 19(4). 10.1097/gme.0b013e31824d8f40

Hyvärinen, M., Juppi, H.-K., Taskinen, S., Karppinen, J. E., Karvinen, S., Tammelin, T. H., Kovanen, V., Aukee, P., Kujala, U. M., Rantalainen, T., Sipilä, S., & Laakkonen, E. K. (2021). Metabolic health, menopause, and physical activity—A 4-year follow-up study. International Journal of Obesity, 46(3), 544–554. 10.1038/s41366-021-01022-x

Janssen, I., Powell, L. H., Crawford, S., Lasley, B., & Sutton-Tyrrell, K. (2008). Menopause and the Metabolic Syndrome: The Study of Women’s Health Across the Nation. Archives of Internal Medicine (1960), 168(14), 1568–1575. 10.1001/archinte.168.14.1568

Juppi, H.-K., Sipilä, S., Cronin, N. J., Karvinen, S., Karppinen, J. E., Tammelin, T. H., Aukee, P., Kovanen, V., Kujala, U. M., & Laakkonen, E. K. (2020). Role of Menopausal Transition and Physical Activity in Loss of Lean and Muscle Mass: A Follow-Up Study in Middle-Aged Finnish Women. Journal of Clinical Medicine, 9(5), 1588. 10.3390/jcm9051588

Juppi, H.-K., Sipilä, S., Fachada, V., Hyvärinen, M., Cronin, N., Aukee, P., Karppinen, J. E., Selänne, H., Kujala, U. M., Kovanen, V., Karvinen, S., & Laakkonen, E. K. (2022). Total and regional body adiposity increases during menopause—Evidence from a follow-up study. Aging Cell, 21(6), e13621. 10.1111/acel.13621

Kambara, T., Ohashi, K., Shibata, R., Ogura, Y., Maruyama, S., Enomoto, T., Uemura, Y., Shimizu, Y., Yuasa, D., Matsuo, K., Miyabe, M., Kataoka, Y., Murohara, T., & Ouchi, N. (2012). CTRP9 Protein Protects against Myocardial Injury following Ischemia-Reperfusion through AMP-activated Protein Kinase (AMPK)-dependent Mechanism. The Journal of Biological Chemistry, 287(23), 18965–18973. 10.1074/jbc.M112.357939

Karvinen, S., Juppi, H.-K., Le, G., Cabelka, C. A., Mader, T. L., Lowe, D. A., & Laakkonen, E. K. (2021). Estradiol deficiency and skeletal muscle apoptosis: Possible contribution of microRNAs. Experimental Gerontology, 147, 111267. 10.1016/j.exger.2021.111267

Kim, H.-R., Rahman, F. U., Kim, K.-S., Kim, E.-K., Cho, S.-M., Lee, K., Moon, O., Seo, Y., Yoon, W.-K., Won, Y.-S., Kang, H., Kim, H.-C., & Nam, K.-H. (2019). Critical Roles of E2F3 in Growth and Musculo-skeletal Phenotype in Mice. International Journal of Medical Sciences, 16(12), 1557–1563. 10.7150/ijms.39068

Klinge, C. M., Riggs, K. A., Wickramasinghe, N. S., Emberts, C. G., McConda, D. B., Barry, P. N., & Magnusen, J. E. (2010). Estrogen receptor alpha 46 is reduced in tamoxifen resistant breast cancer cells and re-expression inhibits cell proliferation and estrogen receptor alpha 66-regulated target gene transcription. Molecular and Cellular Endocrinology, 323(2), 268–276. 10.1016/j.mce.2010.03.013

Korotkevich, G., Sukhov, V., Budin, N., Shpak, B., Artyomov, M. N., & Sergushichev, A. (2016). Fast gene set enrichment analysis [Preprint]. Bioinformatics. 10.1101/060012

Kovanen, V., Aukee, P., Kokko, K., Finni, T., Tarkka, I. M., Tammelin, T., Kujala, U. M., Sipilä, S., & Laakkonen, E. K. (2018). Design and protocol of Estrogenic Regulation of Muscle Apoptosis (ERMA) study with 47 to 55-year-old women’s cohort: Novel results show menopause-related differences in blood count. Menopause, 25(9), 1020–1032. 10.1097/GME.0000000000001117

Krämer, A., Green, J., Pollard, J., & Tugendreich, S. (2014). Causal analysis approaches in Ingenuity Pathway Analysis. Bioinformatics, 30(4), 523–530. 10.1093/bioinformatics/btt703

La Colla, A., Vasconsuelo, A., Milanesi, L., & Pronsato, L. (2017). 17β-Estradiol Protects Skeletal Myoblasts From Apoptosis Through p53, Bcl-2, and FoxO Families. Journal of Cellular Biochemistry, 118(1), 104–115. 10.1002/jcb.25616

Laakkonen, E., Kovanen, V., & Sipilä, S. (2022). Data from Estrogenic Regulation of Muscle Apoptosis (ERMA) study [dataset]. 10.17011/jyx/dataset/83491

Lê Cao, K.-A., Boitard, S., & Besse, P. (2011). Sparse PLS discriminant analysis: Biologically relevant feature selection and graphical displays for multiclass problems. BMC Bioinformatics, 12(1), 253. 10.1186/1471-2105-12-253

Leone, G., DeGregori, J., Yan, Z., Jakoi, L., Ishida, S., Williams, R. S., & Nevins, J. R. (1998). E2F3 activity is regulated during the cell cycle and is required for the induction of S phase. Genes & Development, 12(14), 2120–2130. 10.1101/gad.12.14.2120

Letiges, M., Plomann, M., Standaert, M. L., Bandyopadhyay, G., Sajan, M. P., Kanoh, Y., & Farese, R. V. (2002). Knockout of PKCα Enhances Insulin Signaling Through PI3K. Molecular Endocrinology, 16(4), 847–858. 10.1210/mend.16.4.0809

Leung, K. C., Doyle, N., Ballesteros, M., Sjogren, K., Watts, C. K. W., Low, T. H., Leong, G. M., Ross, R. J. M., & Ho, K. K. Y. (2003). Estrogen inhibits GH signaling by suppressing GH-induced JAK2 phosphorylation, an effect mediated by SOCS-2. Proceedings of the National Academy of Sciences, 100(3), 1016– 1021. 10.1073/pnas.0337600100

Liu, J., Jing, X., Gan, L., & Sun, C. (2012). The JAK2/STAT3 Signal Pathway Regulates the Expression of Genes Related to Skeletal Muscle Development and Energy Metabolism in Mice and Mouse Skeletal Muscle Cells. Bioscience, Biotechnology, and Biochemistry, 76(10), 1866–1870. 10.1271/bbb.120324

Liu, R., Zhou, Z., Zhao, D., & Chen, C. (2011). The Induction of KLF5 Transcription Factor by Progesterone Contributes to Progesterone-Induced Breast Cancer Cell Proliferation and Dedifferentiation. Molecular Endocrinology, 25(7), 1137–1144. 10.1210/me.2010-0497

Love, M. I., Huber, W., & Anders, S. (2014). Moderated estimation of fold change and dispersion for RNA-seq data with DESeq2. Genome Biology, 15(12), 550. 10.1186/s13059-014-0550-8

Martignani, E., Miretti, S., Vincenti, L., & Baratta, M. (2019). Correlation between estrogen plasma level and miRNAs in muscle of Piedmontese cattle. Domestic Animal Endocrinology, 67, 37–41. 10.1016/j.domaniend.2018.12.005

Martone, J., Mariani, D., Desideri, F., & Ballarino, M. (2020). Non-coding RNAs Shaping Muscle. Frontiers in Cell and Developmental Biology, 7, 394. 10.3389/fcell.2019.00394

Matias, M. L., Romao-Veiga, M., Ribeiro, V. R., Nunes, P. R., Gomes, V. J., Devides, A. C., Borges, V. T., Romagnoli, G. G., Peracoli, J. C., & Peracoli, M. T. (2021). Progesterone and vitamin D downregulate the activation of the NLRP1/NLRP3 inflammasomes and TLR4-MyD88-NF-κB pathway in monocytes from pregnant women with preeclampsia. Journal of Reproductive Immunology, 144, 103286. 10.1016/j.jri.2021.103286

Mazur, E. C., Vasquez, Y. M., Li, X., Kommagani, R., Jiang, L., Chen, R., Lanz, R. B., Kovanci, E., Gibbons, W. E., & DeMayo, F. J. (2015). Progesterone Receptor Transcriptome and Cistrome in Decidualized Human Endometrial Stromal Cells. Endocrinology, 156(6), 2239–2253. 10.1210/en.2014-1566

miRBase: The microRNA database. (2022, July 20). https://www.mirbase.org/

Olivieri, F., Ahtiainen, M., Lazzarini, R., Pöllänen, E., Capri, M., Lorenzi, M., Fulgenzi, G., Albertini, M. C., Salvioli, S., Alen, M. J., Kujala, U. M., Borghetti, G., Babini, L., Kaprio, J., Sipilä, S., Franceschi, C., Kovanen, V., & Procopio, A. D. (2014). Hormone replacement therapy enhances IGF-1 signaling in skeletal muscle by diminishing miR-182 and miR-223 expressions: A study on postmenopausal monozygotic twin pairs. Aging Cell, 13(5), 850–861. 10.1111/acel.12245

Pan, J. -l., Yuan, D. -z., Zhao, Y. -b., Nie, L., Lei, Y., Liu, M., Long, Y., Zhang, J. -h., Blok, L. J., Burger, C. W., & Yue, L. -m. (2017). Progesterone-induced miR-133a inhibits the proliferation of endometrial epithelial cells. Acta Physiologica, 219(3), 685–694. 10.1111/apha.12762

Pöllänen, E., Ronkainen, P. H., Suominen, H., Takala, T., Koskinen, S., Puolakka, J., Sipilä, S., & Kovanen, V. (2007). Muscular Transcriptome in Postmenopausal Women With or Without Hormone Replacement. Rejuvenation Research, 10(4), 485–500. 10.1089/rej.2007.0536

Rohart, F., Gautier, B., Singh, A., & Lê Cao, K.-A. (2017). mixOmics: An R package for ‘omics feature selection and multiple data integration. PLOS Computational Biology, 13(11), e1005752. 10.1371/journal.pcbi.1005752

Ronkainen, P. H. A., Pöllänen, E., Alén, M., Pitkänen, R., Puolakka, J., Kujala, U. M., Kaprio, J., Sipilä, S., & Kovanen, V. (2010). Global gene expression profiles in skeletal muscle of monozygotic female twins discordant for hormone replacement therapy. Aging Cell, 9(6), 1098–1110. 10.1111/j.1474-9726.2010.00636.x

Sedano, M. J., Harrison, A. L., Zilaie, M., Das, C., Choudhari, R., Ramos, E., & Gadad, S. S. (2020). Emerging Roles of Estrogen-Regulated Enhancer and Long Non-Coding RNAs. International Journal of Molecular Sciences, 21(10), 3711. 10.3390/ijms21103711

Siles, L., Ninfali, C., Cortés, M., Darling, D. S., & Postigo, A. (2019). ZEB1 protects skeletal muscle from damage and is required for its regeneration. Nature Communications, 10(1), 1364. 10.1038/s41467-019-08983-8

Sipilä, S., Taaffe, D. R., Cheng, S., Puolakka, J., Toivanen, J., & Suominen, H. (2001). Effects of hormone replacement therapy and high-impact physical exercise on skeletal muscle in post-menopausal women: A randomized placebo-controlled study. Clinical Science (London, England: 1979), 101(2), 147–157.

Soneson, C., Love, M. I., & Robinson, M. D. (2016). Differential analyses for RNA-seq: Transcript-level estimates improve gene-level inferences. F1000Research, 4, 1521. 10.12688/f1000research.7563.2

Song, C., Wu, G., Xiang, A., Zhang, Q., Li, W., Yang, G., Shi, X., Sun, S., & Li, X. (2015). Over-expression of miR-125a-5p inhibits proliferation in C2C12 myoblasts by targeting E2F3. Acta Biochimica et Biophysica Sinica, 47(4), 244–249. 10.1093/abbs/gmv006

Sowers, M., Zheng, H., Tomey, K., Karvonen-Gutierrez, C., Jannausch, M., Li, X., Yosef, M., & Symons, J. (2007). 6-year changes in body composition in women at mid-life: Ovarian and chronological aging. The Journal of Clinical Endocrinology & Metabolism, 92(3), 895–901. 10.1210/jc.2006-1393

Statello, L., Guo, C.-J., Chen, L.-L., & Huarte, M. (2021). Gene regulation by long non-coding RNAs and its biological functions. Nature Reviews Molecular Cell Biology, 22(2), 96–118. 10.1038/s41580-020-00315-9

Strzeszewska-Potyrała, A., Staniak, K., Czarnecka-Herok, J., Rafiee, M.-R., Herok, M., Mosieniak, G., Krijgsveld, J., & Sikora, E. (2021). Chromatin-Directed Proteomics Identifies ZNF84 as a p53-Independent Regulator of p21 in Genotoxic Stress Response. Cancers, 13(9), 2115. 10.3390/cancers13092115

Thalacker-Mercer, A. E., Dell’Italia, L. J., Cui, X., Cross, J. M., & Bamman, M. M. (2010). Differential genomic responses in old vs. Young humans despite similar levels of modest muscle damage after resistance loading. Physiological Genomics, 40(3), 141–149. 10.1152/physiolgenomics.00151.2009

Torres, M. J., Kew, K. A., Ryan, T. E., Pennington, E. R., Lin, C.-T., Buddo, K. A., Fix, A. M., Smith, C. A., Gilliam, L. A., Karvinen, S., Lowe, D. A., Spangenburg, E. E., Zeczycki, T. N., Shaikh, S. R., & Neufer, P. D. (2018). 17β-Estradiol Directly Lowers Mitochondrial Membrane Microviscosity and Improves Bioenergetic Function in Skeletal Muscle. Cell Metabolism, 27(1), 167–179.e7. 10.1016/j.cmet.2017.10.003

Uemura, Y., Shibata, R., Ohashi, K., Enomoto, T., Kambara, T., Yamamoto, T., Ogura, Y., Yuasa, D., Joki, Y., Matsuo, K., Miyabe, M., Kataoka, Y., Murohara, T., & Ouchi, N. (2013). Adipose-derived factor CTRP9 attenuates vascular smooth muscle cell proliferation and neointimal formation. The FASEB Journal, 27(1), 25–33. 10.1096/fj.12-213744

Vallejo, G., La Greca, A. D., Tarifa-Reischle, I. C., Mestre-Citrinovitz, A. C., Ballaré, C., Beato, M., & Saragüeta, P. (2014). CDC2 Mediates Progestin Initiated Endometrial Stromal Cell Proliferation: A PR Signaling to Gene Expression Independently of Its Binding to Chromatin. PLoS ONE, 9(5), e97311. 10.1371/journal.pone.0097311

Wang, J., Koganti, P. P., Yao, J., Wei, S., & Cleveland, B. (2017). Comprehensive analysis of lncRNAs and mRNAs in skeletal muscle of rainbow trout (Oncorhynchus mykiss) exposed to estradiol. Scientific Reports, 7(1), 11780. 10.1038/s41598-017-12136-6

Wang, K., Wang, C., Xiao, F., Wang, H., & Wu, Z. (2008). JAK2/STAT2/STAT3 Are Required for Myogenic Differentiation. The Journal of Biological Chemistry, 283(49), 34029–34036. 10.1074/jbc.M803012200

Watts, R., Johnsen, V. L., Shearer, J., & Hittel, D. S. (2013). Myostatin-induced inhibition of the long noncoding RNA Malat1 is associated with decreased myogenesis. American Journal of Physiology-Cell Physiology, 304(10), C995–C1001. 10.1152/ajpcell.00392.2012

Wei, Z., Lei, X., Petersen, P. S., Aja, S., & Wong, G. W. (2014). Targeted deletion of C1q/TNF-related protein 9 increases food intake, decreases insulin sensitivity, and promotes hepatic steatosis in mice. American Journal of Physiology - Endocrinology and Metabolism, 306(7), E779–E790. 10.1152/ajpendo.00593.2013

Winuthayanon, W., Hewitt, S. C., & Korach, K. S. (2014). Uterine Epithelial Cell Estrogen Receptor Alpha-Dependent and -Independent Genomic Profiles That Underlie Estrogen Responses in Mice. Biology of Reproduction, 91(5), 110, 1–10. 10.1095/biolreprod.114.120170

Wu, R.-F., Chen, Z.-X., Zhou, W.-D., Li, Y.-Z., Huang, Z.-X., Lin, D.-C., Ren, L.-L., Chen, Q.-X., & Chen, Q.-H. (2018). High expression of ZEB1 in endometriosis and its role in 17β-estradiol-induced epithelial-mesenchymal transition. International Journal of Clinical and Experimental Pathology, 11(10), 4744–4758.

Zhang, Z., Luo, S., Barbosa, G. O., Bai, M., Kornberg, T. B., & Ma, D. K. (2021). The conserved transmembrane protein TMEM-39 coordinates with COPII to promote collagen secretion and regulate ER stress response. PLOS Genetics, 17(2), e1009317. 10.1371/journal.pgen.1009317

Zhao, L., Wang, J., Li, Y., Song, T., Wu, Y., Fang, S., Bu, D., Li, H., Sun, L., Pei, D., Zheng, Y., Huang, J., Xu, M., Chen, R., Zhao, Y., & He, S. (2020). NONCODEV6: An updated database dedicated to long non-coding RNA annotation in both animals and plants. Nucleic Acids Research, 49(D1), D165–D171. 10.1093/nar/gkaa1046

